# Fighting to persist: Genetic rescue increases long-term fitness despite elevated genetic load

**DOI:** 10.1101/2025.09.06.673525

**Authors:** Jonathan M. Parrett, Mateusz Konczal, Marta Kulczak, Jacek Radwan

**Affiliations:** Evolutionary Biology Group, Faculty of Biology, Adam Mickiewicz University; 60-614 Poznań, Poland

## Abstract

Genetic rescue (GR) can restore fitness in small genetically depauperate populations, however, long-term effectiveness, impact of rescuer quality and introduced deleterious load on population persistence remains uncertain. Using a male-dimorphic mite we tested whether a condition-dependent sexually selected trait influences GR. Genome-wide diversity and genetic load increased similarly for both morphs, while population fitness trajectories differed, suggesting that genetic quality is associated with the trait. Overall, GR sustained fitness benefits over 20 generations. The benefits persisted under thermal stress lowering extinction risk regardless of the rescuer’s morph and introduced load indicating their minimal importance. These results provide experimental evidence that maximising genome-wide variation via GR can yield lasting benefits, even when many deleterious mutations are introduced to populations facing environmental change.

## Introduction

The global rise of human-mediated environmental change and habitat fragmentation poses serious threats to biodiversity (*1–3*), with one of the most immediate concerns being the long-term viability of small, isolated populations (*4, 5*). Such populations are especially vulnerable to declines in genetic diversity due to genetic drift and the subsequent fixation of deleterious mutations, negatively affecting key aspects of population fitness (*6–10*). They may also lack the adaptive capacity and resilience needed under environmental change (*11–13*). Genetic rescue (GR)—the increase in population fitness via introduction of new genetic variants from translocated individuals—offers a potentially powerful conservation strategy to bolster the fitness of imperilled populations (*14–16*) and mitigate the “extinction vortex” (*17*). Numerous studies have shown managed GR or natural migration successfully restoring fitness of inbred populations across diverse taxa (*18–25*). However, uncertainty remains around the persistence of these positive effects (*26, 27*), particularly if a substantial number of deleterious variants are introduced (*28–30*). Indeed, failures of GR to bolster population fitness and even initially positive effects of immigration reversing into population collapse have been reported (*31–34*). If non-genetic drivers of population decline remain, fixed genetic load—both pre-existing and introduced—can reaccumulate and could be particularly severe if populations face environment change which can magnify the negative impacts of deleterious mutations (*35, 36*), potentially reducing fitness and persistence below pre-rescue levels. Alternatively, the relative impact of introduced load may be small compared with the benefits of increasing genetic diversity, which can mask deleterious mutations and ultimately increase population persistence. Given increasing advocacy for GR (*14, 15, 37*), testing how to optimise GR effectiveness and determine the duration of positive effects, most notably in populations facing environmental change, remains an important goal.

Laboratory studies are valuable in this regard, as they enable replicated testing of different GR scenarios across multiple generations, permitting assessment beyond timescales feasible in many natural systems. The results are, however, mixed with some studies demonstrating long-lasting positive effects (*38*), while others showing initial improvements followed by diminished benefits (*39*). Whole-genome resequencing remains underutilised in laboratory GR studies despite its potential in shedding light on these effects. Directly tracking changes in genetic variation over time could clarify wanning GR effects (*39*) and reveal the relative impact of introduced load and genetic variation on changes to fitness and extinction dynamics—which remains disputed (*7, 30, 40*). Characteristics of donor populations and rescuers can also be tested. For instance, the demographic history of donor populations which shapes the type of introduced load can alter GR (*38, 41*). Rescuer sex can also influence GR, with male rescuers providing greater benefits than females (*42*). In polyandrous species males can sire many more offspring compared to what females can produce, increasing the spread of introduced variants more effectively and highlighting GR effectiveness could be impacted by variance in reproductive success of rescuers.

Sexual selection, arising from competition within a sex (typically males), is known to influence important evolutionary processes—including shaping genetic variation and load (*43–45*), consequently affecting adaptation rates and extinction risk (*46–49*)—yet its role in GR has rarely been tested (but see (*39*)). Sexual selection favours traits that increase reproductive success by increasing an individuals’ combativeness or attractiveness (*50*). These sexually selected traits (SSTs) are often condition-dependent, with expression negatively associated with individual genome-wide genetic load (*51*). Thus, variation in SSTs of rescuers could affect the amount of introduced load. Additionally, sexual selection may influence the reproductive success of translocated individuals, where those individuals with high SST expression achieve disproportionate mating success, accelerating GR. Indeed, a recent GR study showed that rescuers from populations evolving under strong sexual selection (*48*) increased GR efficacy, although positive effects were only short-lived (*39*). While initially beneficial, it could be possible that if rescuers dominate reproductive success, subsequent inbreeding may occur earlier between their descendants. Moreover, intra-locus sexual conflict is widespread (*52*) and highly competitive males may increase male-benefit/female-detriment alleles in rescued populations (*53, 54*), potentially constraining population growth as female fitness will generally limit demographic recovery.

Here, using an evolve-and-resequence approach in *Rhizoglyphus robini*, a bulb mite with dimorphic males differing in the expression of a condition-dependent SST, we test whether GR efficacy depends on whether introduced males express a sexually selected weapon (fighters) or lack these armaments (scramblers). Over 20 generations, we assess whether GR improves the fitness and resilience to thermal stress of lines inbred for five generations, leading to an expected inbreeding coefficient of 0.67—a level encountered in many endangered populations which have undergone population bottlenecks (*55*). We examine changes in genome-wide genetic variation resulting from GR, and determine the amount and frequency of initially fixed and introduced deleterious mutations prior to the thermal extinction assay. These patterns are linked to changes in fitness and extinction risk of rescued populations.

### Changing genomic variation due to genetic rescue

Genetic variation (nucleotide diversity: π) was calculated from pool-seq samples of 18 lines that had underwent five generations of full sib mating (hereafter; inbred lines) and at three time points after GR from either two fighter or scrambler males at F_2_, F_6_ & F_10_ (Fig. 1a). Genome-wide π was used to estimate changes in genetic diversity quantified as the ratio of genetic variation of evolved populations with GR at each generation to inbred line genetic variation (Δπ). Most populations had values of Δπ greater than 1 at F_10_ (Δπ: 95% CI 1.29 – 1.57; Fig. 1b), demonstrating GR was generally successful in restoring genetic variation. There was no difference in Δπ between lines rescued with fighter or scrambler males (*χ^2^* = 0.26, *df* = 1, *p* = 0.609), but Δπ did increase over the course of the experiment (*χ^2^* = 9.00, *df* = 1, *p* = 0.003; Fig. 1b; Table S1). Similar results were found when estimating genetic variation by the number of segregating sites (Watterson’s ϴ: fig S1; Table S2).

**Fig. 1.**
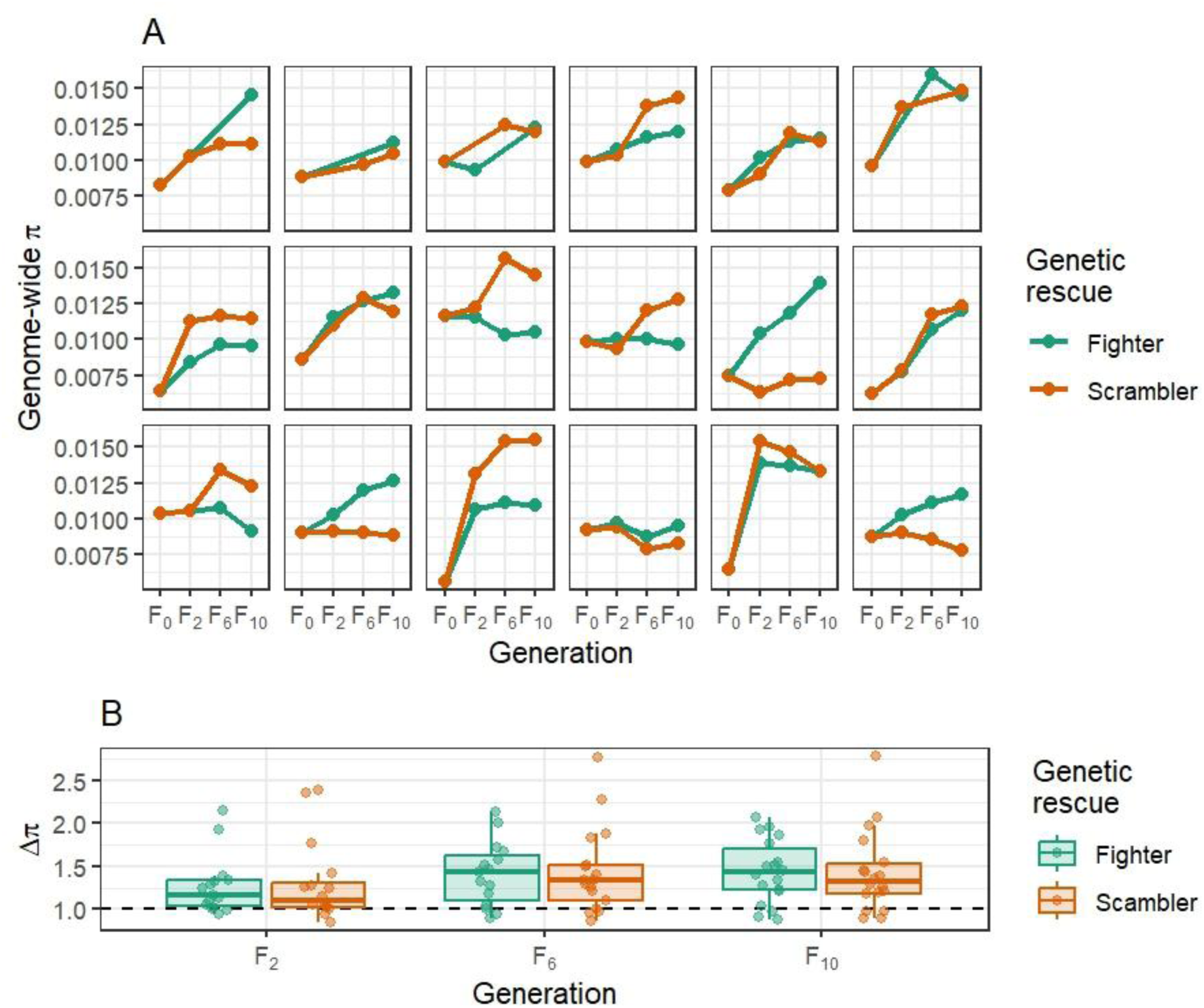
Changes in genome-wide nucleotide diversity (π) due to GR in F_2_, F_6_ and F_10_. **(A)** shows mean genome-wide estimated π from 10kb sliding windows (with 10kb steps) in inbred lines (F_0_) and subsequent generations post GR (F_2_, F_6_ & F_10_). **(B)** represents estimated changes in genome-wide π at each generation post GR after controlling for inbred line measurements (Δπ = π at F_2_ (F_6_ & F_10_) / inbred line π). Thus, a value of 1 for Δπ (dashed horizonal line) would indicate no change in genome-wide variation, a value less than 1 indicating loss of genetic variation and a value greater than 1 indicating an increase. In both A and B GR populations with rescue from fighters and scramblers are shown in green and orange, respectively. In B boxes are composed of the median and hinge values (25th and 75th percentiles), with whiskers ± interquartile range × 1.5. Individual data points from each population denoted by circles.

Deleterious mutations are expected to be under mutation-selection balance (*56, 57*), and we identified presumed deleterious mutations in inbred lines and the evolved GR populations (F_10_) based on their frequency in the stock population from which inbred lines and rescuers were derived. Specifically, non-synonymous missense and nonsense mutations segregating in the stock population with a frequency < 0.1 were assumed to be enriched in deleterious variants (hereafter; deleterious mutations). As genome-wide variation declines, due to reduced Ne and inbreeding, it is expected that an increasing number of variants will become fixed in populations (*6, 8*). Indeed, there was a negative correlation between fixed number of missense and nonsense deleterious mutations in inbred lines and genome-wide π (*r* = −0.77 and *r* = −0.76, respectively)—which were themselves non-randomly distributed across chromosomes within each inbred line (fig. S2).

As deleterious mutations are expected to be (partially) recessive, introduction of alternative variants would be predicted to mask a considerable amount of fixed load in offspring sired by rescuers and heterozygotes in subsequent generations (*26, 27*). Despite this benefit from GR in restoring heterozygosity and masking fixed load, some concern remains with the amount of load being introduced to populations via GR (*30*). By quantifying the frequency of deleterious mutations that were either initially fixed in inbred lines or introduced via GR we explored the fate of such alleles at F_10_ (Fig 2). As would be expected there was a positive association between GR effectiveness in restoring genome-wide diversity (Δπ) and the amount of deleterious mutations introduced at F_10_, whether estimated by number (fig S3; Table S3; missense: *χ^2^* = 13.62, *df* = 1, *p* < 0.001; nonsense: *χ^2^* = 16.75, *df* = 1, *p* < 0.001) or their sum of frequencies (fig S4; Table S4 missense: *χ^2^* = 45.94, *df* = 1, *p* < 0.001; nonsense: *χ^2^* = 47.57, *df* = 1, *p* < 0.001). Furthermore, those chromosomes most burdened with fixed deleterious load (standardised by chromosome length) had on average the greatest values of Δπ at F_10_ (*χ^2^* = 71.2, *df* = 1, *p* < 0.001; fig. S5; Table S5) and within individual replicate GR populations correlation coefficients between per-chromosome fixed deleterious load and per-chromosome Δπ were generally positive (95% CI 0.23 – 0.50; fig. S6)—suggesting that selection after GR favoured the introduction of new variants in regions of the genome more burdened with fixed load.

**Fig 2.**
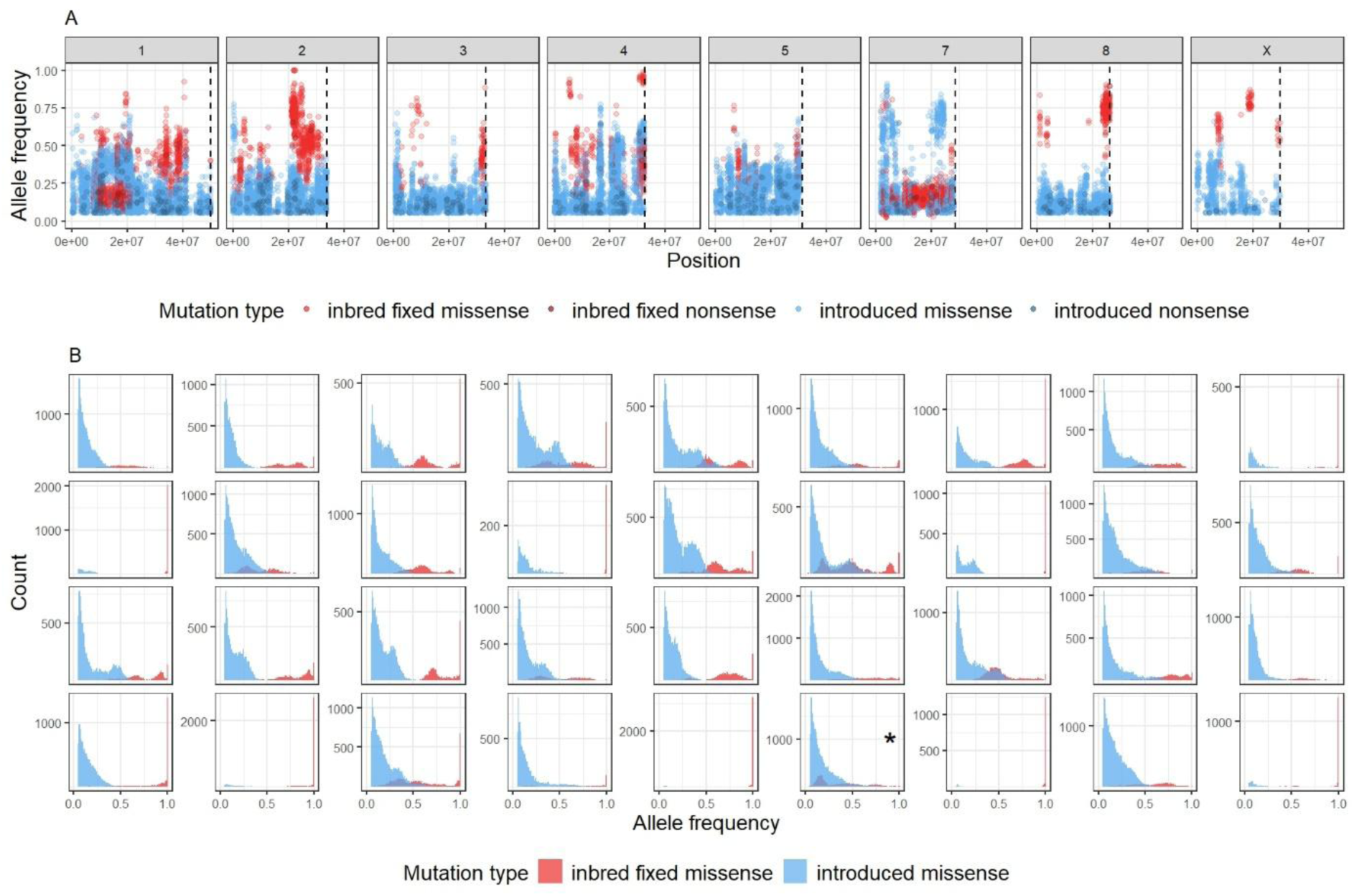
Frequencies of initially fixed and introduced deleterious mutations at F_10_. **(A)** provides an example from a single replicate of allele frequencies of deleterious missense (lighter shading) and nonsense (darker shading) mutations that were initially fixed in the inbred line (red) and introduced via GR (blue) at F_10_ across each chromosome. Vertical dashed lines represent the length of each chromosome. **(B)** summary histograms of all replicates with GR showing distribution of deleterious missense mutations initially fixed in inbred lines (red) and introduced via GR (blue) at F_10_ from across all entire genomes. Histograms in the two top rows are replicates with GR from fighter males and the two bottom rows are replicates with GR from scrambler males. The plot highlighted with an asterisk (*) is the example replicate from A. For the corresponding summary histograms of nonsense mutations see fig. S7.

The mean frequency of initially fixed deleterious mutations at F_10_ ranged from being close to 1 through to 0.3, representing ineffective and effective selection against initially fixed load in inbred lines because of GR, respectively. We find no indication that either morph was more effective in reducing the frequency of initially fixed inbred load, measured as the mean frequency of initially fixed deleterious mutations at F_10_ (missense: *χ^2^* = 0.57, *df* = 1, *p* = 0.452; nonsense: *χ^2^* = 0.33, *df* = 1, *p* = 0.566; Fig. 3a). GR was less effective in reducing the frequency of initially fixed load on the sex chromosome compared to autosomes (missense: *χ^2^* = 25.67, *df* = 1, *p* < 0.001; nonsense: *χ^2^* = 13.30, *df* = 1, *p* < 0.001; Fig. 3b) likely due to rescuers used here being the heterogametic sex (*43*).

**Fig. 3.**
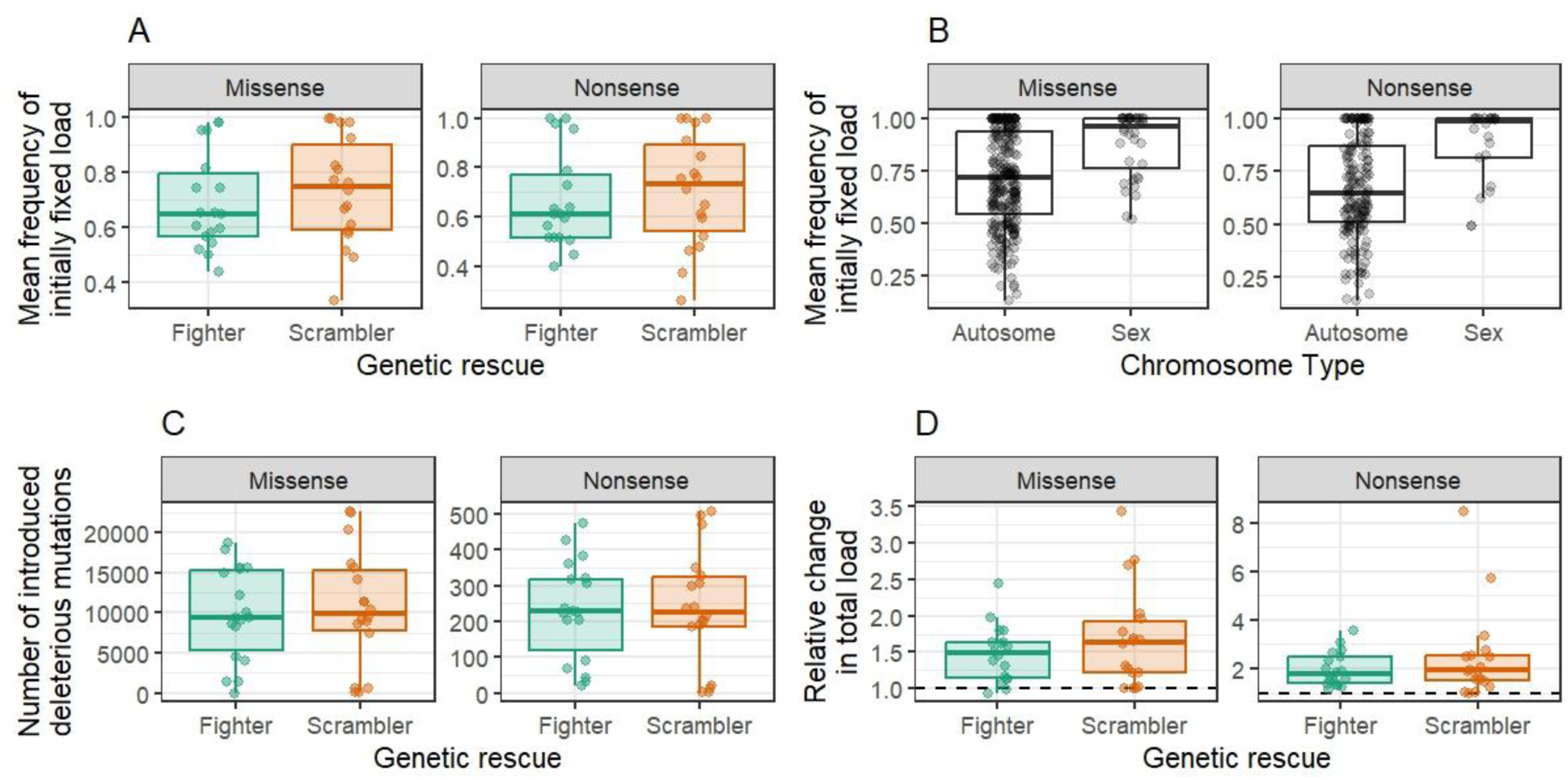
Changing deleterious load in populations after GR. **(A)** the mean frequency of missense and nonsense mutations at F_10_ that were initially fixed in inbred lines when inbred lines had GR from either fighter (green) or scrambler (orange) males. **(B)** the mean frequency of missense and nonsense mutations at F_10_ that were initially fixed in inbred lines found on autosomes compared to those found on the sex chromosome. **(C)** total number of missense and nonsense mutations at F_10_ introduced to populations via GR from fighter (green) or scrambler (orange) males. **(D)** change in total missense and nonsense load segregating in populations due to GR from either fighter (green) or scrambler (orange) males at F_10_ was generally positive (missense: 95% CI 1.42 – 1.81; nonsense: 95% CI 1.78 – 2.78). Calculated as the sum of allele frequencies of both initially fixed missense (or nonsense) and introduced load at F_10_ divided by the number of initially fixed deleterious missense or nonsense mutations, where a value of 1 would indicate no change in load (horizonal dashed lines), values greater than 1 an increase in total load and values less than 1 a decrease in total load. Boxes are composed of the median and hinge values (25th and 75th percentiles), with whiskers ± interquartile range × 1.5. With individual data points in A, C and D showing individual replicates and in B every chromosome from each replicate.

The total number of introduced deleterious mutations often far exceeded those that were initially fixed in inbred lines (Fig. 2b & fig. S7). We, however, find no support that fighters, previously shown to have fewer deleterious mutations and selecting against them more strongly (*43, 58*), introduce fewer deleterious mutations than scramblers whether controlling for Δπ or not (Fig. 3c; Table S3). The increase in mean frequency of introduced deleterious mutations were on average less than the comparative decrease in mean frequency of fixed inbred deleterious mutations (missense: *χ^2^* = 15.49, *df* = 1, *p* < 0.001; nonsense: *χ^2^* = 18.08, *df* = 1, *p* < 0.001; Fig. 2b and fig. S7). This pattern likely reflects that two males were used in GR, or four introduced genomes each contributing to reducing the frequency of initially fixed mutations but carrying their own private deleterious mutations that remained at comparatively low frequency. Nevertheless, many of the introduced deleterious mutations reached frequencies greater than that they segregated at within the stock population. Consequently, GR generally increased the total load segregating in populations (Fig. 3d), with no evidence that either male morph led to higher overall load in populations (missense: *χ^2^*= 1.671, *df* = 1, *p* = 0.196; nonsense: *χ^2^* = 1.26, *df* = 1, *p* = 0.261). If an inbred line represented a random genome from a source population, GR should on average mask the same number of deleterious mutations as it introduces. That GR generally increased total genetic load and increases in nonsense load was comparatively higher to missense load may reflect purging of strongly deleterious variants when establishing inbred lines.

### Fitness effects of genetic rescue

The fecundity of male and female pairs was determined by a significant two-way interaction between GR treatment and generation (*χ^2^* = 30.65, *df* = 6, *p* < 0.001; Fig. 4a, Table S6). Importantly, and irrespective of the morph of rescuer males, GR significantly increased fecundity when compared to inbred lines without GR, with this positive effect of GR maintained across 10 generations— demonstrating the long-term effectiveness of GR in bolstering population fitness. Interestingly, the effect of GR differed over time depending on the male morph used as rescuers. In the first two generations post GR the increase in fecundity was significantly greater when scrambler males were used for GR compared to fighters, however, in the latter half of the experiment the fecundity of pairs from fighter rescued inbred lines had significantly greater fecundity compared to those rescued by scramblers (Fig. 4a, Table S6). The mean fecundity of population in F_10_ was a good predictor of total population productivity in F_11_ (*χ^2^* = 61.51, *df* = 2, *p* < 0.001; fig S8)—supporting these measures of fecundity to be a good estimate for overall population fitness and growth potential.

**Fig. 4.**
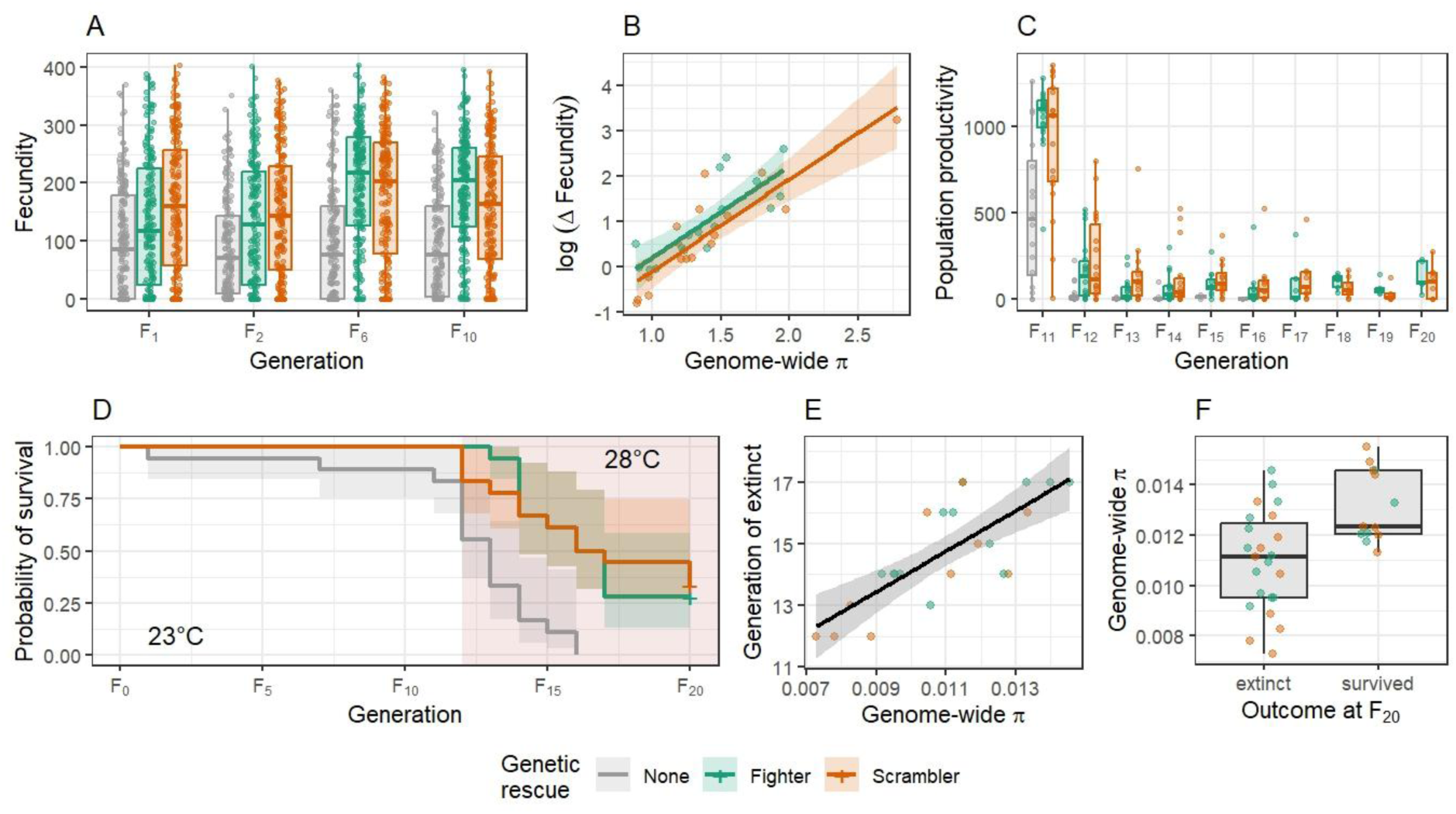
Fitness consequences of GR and associations with nucleotide diversity (π). **(A)** Fecundity of male and female pairs at generations F_1_, F_2_, F_6_ and F_10_ from nine days of egg laying. **(B)** Association between changes in genome-wide nucleotide diversity (Δπ) and change in log transformed mean fecundity (Δfecundity) of individuals from populations with GR at F_10_. **(C)** Productivity of populations from F_11_ to F_20_ representing all adults and tritonymphs from six days of oviposition. Population productivity in F_11_ was carried out at 23°C, all other generations F_12_ to F_20_ populations were reared at 28°C. **(D)** Proportion of populations surviving across the entire experiment from F_0_ to F_11_ when reared at 23°C and then at 28°C from F_12_ to F_20_, the increase in temperature is denoted by red shading. **(E)** Association between absolute genome-wide π at F_10_ and generation of extinction of populations with GR. **(F)** Comparison of absolute genome-wide π at F_10_ of populations with GR that either went extinct during the thermal extinction assay or survived until F_20_ when the experiment ended. In all plots populations with GR from fighter and scrambler males are shown in green and orange, respectively. In A, C and D those inbred lines without GR are shown in grey. Note that in E and F points are coloured for clarity but there was no significant difference between GR treatments. In A, C and F boxes are composed of the median and hinge values (25th and 75th percentiles), with whiskers ± interquartile range × 1.5. In B, D and E shaded areas indicate 95% CIs.

Although there was no significant difference in changes between segregating load when GR came from fighter or scrambler males (Fig. 3d), the significant morph effect on fecundity that interacted across generations (Fig. 4a) taken together with Δπ generally increasing across the experiment (Fig. 1b), suggests that as introduced variants increase in frequency their effect on population fitness differs and depends on whether GR came from fighter or scrambler males. This interaction could be the result of fewer strongly sexually antagonistic variants introduced by scrambler rescuers (*53, 59*) giving scrambler-rescued populations a short-term advantage, but as introduced variants increased in frequency and more frequently became exposed to selection, the longer-term benefits may be mostly associated with higher genetic quality of fighters (*43, 58*). As we are unable to determine the nature and strength of deleterious mutations in these data our proxy for deleterious mutations may lack the power to explain these subtle fitness effects.

Notwithstanding the effect rescuer morph had on GR, it generally increased inbred line fitness. There were, however, obvious idiosyncratic effects of whether GR led to increases in fecundity and several attempts failed. The significant association between Δπ and changes in mean fecundity (Δfecundity: mean GR rescued fecundity / mean inbred line fecundity) of populations at F_10_ (*χ^2^* = 50.44, *df* = 1, *p* < 0.001; Fig. 4b; Table S7) indicates that the failure of GR to bolster population fitness is likely a consequence of introduced males or their descendants having little to no reproductive success and subsequently no or very few genetic variants being introduced. This increase in fitness can likely be attributed to masking the negative effects of initially fixed deleterious mutations in heterozygotes, and we ran models to examine if the frequency of initially fixed missense and nonsense at F_10_ could explain any additional variance in Δfecundity compared to Δπ alone. Indeed, the mean frequency of initially fixed deleterious missense and nonsense mutations after GR at F_10_ were strong individual predictors of Δfecundity (missense: *χ^2^* = 29.69, *df* = 1, *p* < 0.001; nonsense: *χ^2^* = 23.62, *df* = 1, *p* < 0.001), however, these significant effects disappeared when changes in genome-wide diversity (Δπ) were also included in the models. This indicates that Δπ is likely a good proxy for the potential benefits of masking of deleterious mutations in small populations during GR. Importantly, the negative effects of introduced deleterious mutations will on average be masked when segregating at low frequency (Fig. 2b; fig S7) due to the majority being (partially) recessive. Coupled with masking of initially fixed deleterious load this can explain why despite GR increasing total load in populations it still substantially increases population fitness (Fig. 4a). Moreover, some of this variance in efficacy of GR in restoring population fitness can be attributed to inbred line π—the most genetically depauperate inbred lines, with lowest fecundity (fig. S9), benefitted the most for GR and had the greatest relative increase in fecundity at F_10_ (fig. S10; Table S8). There was, however, no significant effect of rescuer morph or recipient population morph proportion on these relationships (Table S8). Previously it was shown using outbred individuals that scramblers suffer low fitness when their rivals are fighters (*60, 61*). The lack of difference between male morphs here suggests that even less reproductively competitive males, without SSTs, can still achieve relatively high reproductive success compared to resident inbred males. Whether this result would hold if using inbred rescuers (*41*) or in taxa without discrete SSTs and mating strategies remains largely unknown.

These results suggest that any effect of introduced load in affecting fitness is small, likely due to those positions remaining heterozygous in populations, but given enough time with populations maintained with low Ne or after subsequent bottlenecks introduced load could fix and negatively impact populations. We explored the latter scenario, by exposing populations to thermal stress to test whether GR increases or decreases population persistence and resilience to simulated climate change. There was significant decline in all population’s productivity following an increase in temperature of 5°C between F_11_ and F_12_ (*χ^2^* = 34.89, *df* = 1, *p* < 0.001; Fig. 4c; Table S9), with those inbred lines without GR having lower overall productivity compared to those with GR (*χ^2^* = 11.26, *df* = 2, *p* = 0.004; Fig. 4c). There was also a general decline in population productivity of surviving lines in further generations (F_12_ to F_20_) after the temperature was increased (*χ^2^*= 6.30, *df* = 1, *p* = 0.012), again with those inbred lines without GR having significantly lower overall productivity compared to GR populations (*χ^2^* = 22.69, *df* = 2, *p* < 0.001; Fig. 4c; Table S10). These patterns are also reflected in population survival across the entire experiment with GR having a positive effect on persistence (*χ^2^* = 20.94, *df* = 2, *p* < 0.001; Fig. 4d; Table S11). Extinctions occurred in three inbred lines without GR under benign temperatures and greatly increased in frequency once temperature was increased, with all inbred lines without GR being driven to extinction by F_16_. Contrastingly, no lines with GR became extinct in benign conditions prior to increases in temperature. Thermal stress also drove some GR populations to extinction, but the rate was significantly less compared to those without GR and in total 11 (5 & 6 rescued by fighter and scrambler males, respectively) survived until F_20_ when the experiment was ended. Notably, resilience and extinctions rates did not significantly differ between populations rescued by fighters or scrambler males (Fig. 4d; Table S11).

Absolute genome-wide variation of populations with GR (π in F_10_) was associated with their persistence under thermal stress. Of those that went extinct, there was a positive association between generation of extinction and genome-wide variation (*χ^2^* = 22.28, *df* = 1, *p* < 0.001; Fig. 4e; Table S12). Ultimately, those lines with GR that survived until the end of the experiment at F_20_ had significantly higher genome-wide variation at F_10_ compared to those that went extinct during the thermal extinction assay (*χ^2^* = 10.82, *df* = 1, *p* < 0.001; Fig. 4f; Table S13). Indicating that introduced load during population bottlenecks unlikely contributed to increasing extinction risk. Overall, these results demonstrate little risks associated with introduced deleterious mutations through GR and suggest that maximising genome-wide variation, as suggested by others (*7, 40*), to be a priority and could buy time needed for other interventions to be implicated in small, imperilled populations facing environmental change.

## Methods

### Model organism: Rhizoglyphus robini

The bulb mite, *Rhizoglyphus robini*, is a widespread species commonly found in and on the bulbs of various plant species, including several of agricultural and economic importance. There is a growing body of literature and research using *R. robini* as a model organism for research in ecology and evolution, most notably associated to sexual selection (*43, 46, 62*) and the expression and maintenance of alternative male reproductive tactics (*53, 63–65*). The molecular resources available for this species are increasing and improving with several studies of transcriptomics and genomics (*43, 59, 66*), including a recently published chromosome-level assembly and recombination landscape (*67*).

Male *R. robini* exhibit two distinct reproductive strategies: fighters and scramblers. Fighters possess a thickened and terminally pointed third pair of legs, which they use in aggressive contests with rival males. In contrast, scramblers have legs of approximately uniform thickness and avoid direct competition, instead adopting a ‘sneaky’ reproductive strategy (*68*). Male–male fights can occasionally escalate into lethal battles, in which scramblers experience higher mortality than fighters (*61*). Moreover, when populations are biased toward fighters, scrambler males inseminate fewer females and sire fewer offspring than fighter males likely due to higher mortality (*60*). In addition to the competitive social environment, abiotic factors such as temperature can modulate the fitness of each male morph (*69*), however, short-term acute thermal stress appears to negatively affect morph fitness in similar ways (*70*). Morph determination in this species is complex, being both heritable with considerable additive genetic variance (*71, 72*) and environmentally determined. For example, fighter expression is also condition-dependent and there is an increased probability of larger tritonymphs (final juvenile stage) developing into fighter males, and decreasing food availability during development increases the probability of males developing into scramblers (*63, 71, 73*). Fighter males have also been shown to have fewer deleterious recessives compared to scramblers (*58*) and that populations artificially selected for fighter expression increases the strength of purging of presumed deleterious mutations across much of the genome (*43*). Despite the benefits of purging deleterious mutations through male-male competition, females from fighter selected populations have reduced fecundity (*53*) likely because of sexually antagonistic pleiotropic effects of fighter beneficial alleles being expressed in females (*59*), which can increase risk of extinction under thermal stress (*74*). Morph expression of males (*72*) and female fecundity (*75*) are both sensitive to inbreeding in patterns suggesting polygenic genetic underpinnings, however, the former may also be associated with a (few) large effect loci (*72*).

### General husbandry and stock population

An outbred stock population of *Rhizoglyphus robini* was established from ca. 200 individuals collected in 2017 from onions found in fields near Mosina, Poland. The stock population was housed in a plastic container (∼ 7 x 10 cm) with small holes punched into the bottom, experimental lines were housed in round plastic containers (∼ 2 cm Ø) and individuals or pairs in glass vials (∼ 1cm Ø). All containers had a plaster of Paris base (∼ 1 cm) that was soaked in water and placed on wet tissue paper to maintain high humidity (>90%). During the maintenance of the stock population its size was kept large (> 10,000 individuals) and eggs collected and transferred to new containers roughly twice a month. Mites were fed *ad libitum* with powdered yeast 2 or 3 times a week dependent on life stage, with water sprayed onto tissue paper to ensure adequate humidity. All mites were kept in a dark incubator at 23°C (unless stated otherwise). All populations and lines were kept within larger plastic boxes and surrounded by water and washing liquid to ensure contamination from other sources could not occur. All manipulation of mites was carried out by carefully moving them with fine needles under dissecting microscope, similarly any counting of eggs or individuals were performed under dissecting microscope at adequate magnification.

### Establishing inbred lines and Genetic rescue (GR) protocol

The stock population was synchronised by life stage by collecting eggs and allowing individuals to develop. During the last juvenile stage (tritonymph) ∼200 individuals were isolated in vials and upon maturation into adults their sex determined, thus all individuals were guaranteed to be unmated. 40 inbred lines were established by pairing a single male and female haphazardly in plastic containers, pairs were allowed to mate and produce eggs over the next three days, after which the male and female were removed. Seven days after removing adults, 12 nymphs (or as many as possible) were individually isolated into vials and five days later individuals were sexed. From each inbred line two brother and sister pairs were made. From each inbred line, the total number of adults produced by a single male and female was determined by counting and removing adults from plastic containers every other day, three times to ensure all were counted. These counts, plus those from vials, were used to track the effect of inbreeding on inbred line productivity.

Following the same timeline as above, one of the containers from each inbred line was randomly selected and checked for nymphs. If there were adequate numbers of nymphs (e.g. 12 or more) they were isolated into vials. If the first container did not have 12 nymphs the second container was inspected, if there were at least 12 nymphs they were isolated, and if both containers did not have 12 nymphs, then one of them was randomly selected to isolate nymphs from. The total number of adults produced by each pair were counted as above. This protocol was repeated a further four times, thus each inbred line used in the further GR experiment was established from a total of five generations of full sib x sib matings. On occasion we did not have enough males and therefore males from the corresponding plastic container of that inbred line were used, females from plastic containers were never used. There was a clear decline in number of adults produced by male and female pairs as the number of generations of inbreeding increased (*χ^2^* = 73.56, *df* = 1, *p* < 0.001; fig S11) and during the inbreeding protocol seven inbred lines were lost due to both male and female pairs not producing any offspring (fig S11). Surviving inbred lines were then allowed to expand for ca. 5 generations before the GR experiment began. During this period within an inbred line individuals could freely mate and reproduce. The size of population of more productive inbred lines were regulated by removing approximately half of the population (e.g. all life stages) once a week, in less productive inbred lines this regulation of population size was carried out less frequently. During this period of population expansion, a further two inbred lines were lost. Then from each inbred line we then sampled for genomic material (see below) and seven days later transferred 50 females (or as many as possible) to new containers to lay eggs for six days, which upon development and maturation were used in the full GR experiment. Over this egg laying period, we also collected eggs from the stock population to synchronise life stages with the inbred lines and therefore ensure all adults used were all approximately the same age.

From the 30 surviving inbred lines, we randomly selected 18 to be used in the full experiment. From each inbred line a control replicate was established and consisted of 20 females and 20 males transferred to a round plastic container. At the same time a further two replicates were established consisting of 20 females and 18 males, in these two replicates we introduced either 2 fighter or 2 scrambler males from the stock population. The morph of all males used were recorded. We consider the establishment of these replicate populations as F_0_ of the experiment. Three days later all individuals were transferred to a second container for a further three days of egg laying before all adults were discarded. The second container was used for replicate maintenance and fecundity assays (see below), unless it did not contain enough individuals and then we also included individuals from the first container. Four days after the adults were discarded and to standardise the density of replicates, we transferred approximately 100 larvae (or as many as available) to new containers, which ten days afterwards, roughly a day after most individuals had eclosed as adults, we haphazardly transferred 20 females and 20 males to new containers for egg laying and the production of the next generation as described above. If there were not enough adults, nymphs were also transferred to make the total number of individuals up to 40. We recorded the generation when a replicate became extinct and was determined by there either being a) no individuals or b) the only adults were the same sex. This protocol was repeated for a further ten generations before the thermal extinction assay began (see below).

### Fecundity and population productivity assays

Periodically throughout the experiment we assayed the fecundity of male and female pairs, these fecundity assays were performed at F_1_, F_2_, F_6_ and F_10_ (where F_0_ is the establishment of replicates without GR or with GR from fighter or scrambler males). From each replicate population we isolated 30 larvae (or nymphs) in vials, which upon their maturation as adults were sexed, all adult mites used were < 3 days old. From each replicate we attempted to establish twelve male and female pairs, which were transferred into new vials. Only individually isolated females were used, but on the rare occasion there were not enough available males they were taken from containers. Pairs were housed for five days, before being transferred to another new vial for a further four days of egg laying before being removed. Immediately after adults were removed from the vials they were frozen at −20°C for egg counting at a later date. The fecundity assay therefore represents nine days of potential egg laying, with the consideration that females take approximately a day to start laying eggs after their first mating. Any deaths were recorded and if the male was found dead in the first vial a replacement male was used. From each vial the total number of eggs (and on occasion larvae) were counted. Preliminary testing found that eggs did not break during the freezing and thawing and therefore the freezing of vials will not influence our measurements of fecundity.

At F_11_ we also carried out a total population productivity assay, which consisted of counting all adults and tritonymphs in the first and second egg laying containers, plus those in the new container in which ∼100 larvae were transferred during replicate maintenance (e.g. total productivity from six days of egg laying). The containers were frozen 14 days after adults were removed, following which the entire contents of containers were washed over fine nets to remove any remaining yeast. The dead mites were then spread across a small piece of squared paper and counted.

### Thermal extinction assay

In addition to tracking population level effects of GR in benign conditions (described above), we also tested whether GR increased population resilience to thermal stress and an increase of 5°C. After F_11_ adults had been removed from egg laying containers, the F_12_ eggs were then placed in an incubator at 28°C and all replicates maintained at this increased temperature for the remainder of the experiment. The protocol remained the same as above, except that the development times were faster at 28°C compared to 23°C: instead of 14 days from removing adults (F_n_) in second containers to transferring 40 F_n+1_ adults to egg laying containers this period was reduced to 11 days. As above, a replicate was considered extinct if there were a) no individuals or b) the only adults were the same sex. In addition, we tracked population productivity each generation in the same way as described above by freezing and counting adults and nymphs from all containers.

### Genomic sampling, mapping and analysis

For genomic analyses we sampled material at several time points and carried out whole-genome resequencing on pools of individuals. During the expansion of inbred lines prior to GR events we sampled 20 females from each (n = 18), further from those replicates with GR from either fighter (n = 18) or scrambler (n = 18) males we sampled 40 females from three generations (F_2_, F_6_ and F_10_)— giving a total of n = 126 samples for resequencing. Adult females were randomly selected from inbred lines or after population maintenance in those with GR. All samples were placed in ATL buffer before freezing at −20°C. The tissue of all individuals was carefully homogenised using microtube pestles and DNA extracted by Proteinase K digestion (24 h) followed by standard procedures using QIAamp UCP Micro Kit (Qiagen). DNA quality and concentration was assessed using agrose gels and Qubit double-stranded DNA HS Assay Kit. Several samples were flagged as likely being degraded and/or having low concentration, however, as no additional samples were collected, we decided to proceed with resequencing.

Library preparation and whole-genome resequencing was performed by Novogene (UK) using the Illumina Nova-Seq X platform with 10B flow cell to produce 2 ×150 bp reads (average 241.7 × 10^6^: range: 170.5 × 10^6^ – 330.3 × 10^6^). Adaptors were trimmed from reads using Trimmomatic (v.0.39) (*76*) and any unpaired reads discarded. Fastq files were mapped to the reference genome assembly (*67*) with bwa mem (v.0.7.17-r1188) (*77*) using default settings. Sam files were converted to bam files, sorted, duplicates marked, and ambiguously mapped reads discarded using samtools (v.1.9) (*78*). There was considerable variation in the percentage of reads from each sample that were mapped successfully (range 23% - 94%), the average overall percentage of reads that were mapped from each sample was 72%, of which an average of 15% (range 13 −19%) were removed to being marked as duplicates or ambiguously mapped. After cleaning reads, we were left with an average of 148.5 × 10^6^ reads per sample, ranging from 39.0 × 10^6^ – 228.6 × 10^6^.

Using samtools we created an mpileup file excluding positions with a quality score < 30. Next, we extracted the coverage of each position in 10kb steps to draw a distribution of expected coverage across the genome for each sample. As a consequence of the considerable variation in expected coverage, we decided to exclude nine samples from further analysis as they would have a large impact on subsequent subsampling steps. We retained all inbred line samples and F_10_ samples with five F_2_ and four F_6_ samples being excluded (6 and 3 rescued from fighter and scrambler males, respectively). Hereafter, we refer to the inbred line and three possible time points after GR from both fighter or scrambler males of that inbred line as an experimental block to denote larger scale hierarchy in experimental design, this does not refer to any temporal difference within the experiment. To retain as much data as possible we opted to perform filtering and subsampling of samples within individual experimental blocks. Thus, all comparisons between inbred line samples and any evolved line derived from the same inbred line were based on the same filtering criteria, but the same filtering thresholds are not necessarily the same between experimental blocks. Next, to minimise calling false SNPs using PoPoolation2 (*79*) we identified indels using identify-genomic-indel-regions.pl and filtered indels (plus 5 bp either side), based on 5% of total expected coverage within an experimental block using filter-pileup-by-gtf.pl. Furthermore, repeats identified by (*67*) were filtered, again using filter-pileup-by-gtf.pl.

Next, using PoPoolation (*80*) we estimated genomic variation either as nucleotide diversity (Tajima’s Pi: π) or the number of segregating sites (Watterson’s theta: ϴ) in 10kb sliding windows. First, we subsampled (without replacement) the mpileup file of each sample using subsample-pileup.pl. Within each experimental block we visualised the expected coverage of all samples to determine subsampling parameters, which were approximately 50% of the mean expected coverage of all samples within an experimental block (median 35×: range 25× - 50×). Then using Variance-sliding.pl we estimated π and ϴ in 10kb sliding windows, using a 10kb step size, a pool size of 40 for inbred samples and 80 for F_2_, F_6_ and F_10_ samples with GR, and a minimum count of 3 or 5% (which ever was largest) for a SNP to be called, a phred score of >30 and any window with < 20% coverage was discarded. Within each experimental block all windows from each sample had to meet our criteria, otherwise they were discarded.

Finally, in addition to the above resequencing we also sampled and resequenced 200 females from the stock population, split between two pools of 100 individuals. All procedures were identical to those described above, except that DNA extractions were performed using a DNeasy Blood and Tissue Kit (Qiagen). 150bp × 2 reads obtained from an Illumina Nova-Seq X platform had adaptors trimmed, mapped to the reference assembly, followed by filtering indels and repeats (as described above), with the final number of mapped reads in each sample after removing duplicates and ambiguously mapped reads being 170.7 × 10^6^ and 181.2 × 10^6^. These stock population samples were used to identify presumed deleterious mutations segregating in the population, which were fixed in inbred lines or introduced to a population via GR. Specifically, we identify non-synonymous positions in coding regions of genes that were expressed at a mean level of fragments per kilobase of transcript per million mapped reads (FKPM) >1 (*66*) and where the minor allele had a frequency < 0.1. We assume that mutations causing non-synonymous amino acid change (missense) or a stop codon (nonsense) and segregating at low frequencies in the stock population are, on average, likely deleterious in nature, with the latter assumed to be more strongly deleterious.

To achieve this, an mpileup file in VCF format was created from the bam files of the stock population, inbred line and F_10_ samples using bcftools (v1.11) (*81*), with quality scores >30 and retaining only coding regions of expressed genes by filtering using a bedfile (made using bedtools v.2.27.1 (*82*)) of genes with an FKPM > 1. The VCF file was then annotated using SnpEff (v5.2) (*83*) to identify non-synonymous positions. First, we filtered the annotated VCF file to non-synonymous positions only and to where the minor allele was < 0.1 in the stock samples, further filtering based on coverage was performed and all positions had to have > 50× from stock population samples (sum of both) and > 25× in all inbred and F_10_ samples. For each inbred line sample, we then only retained positions in which the presumed deleterious mutations were completely fixed and represent inbred fixed load: note that this filtering procedure also retained SNPs that were not detected in the stock population sample likely due to their very low frequency. Next, we checked the frequency of all these positions in F_10_ samples that had been determined as fixed load in inbred lines. As most deleterious mutations are expected to be recessive or partially recessive an increase of frequency of the major allele is assumed to reflect the potential masking effects of fixed inbred load via GR. Finally, we quantified the amount of presumed deleterious alleles introduced to populations as a consequence of GR by filtering files to only monomorphic positions in inbred line samples where major alleles (in the stock population) were fixed, followed by determining the frequency of any previously identified presumed deleterious mutations identified in the stock population in F_10_ samples derived from the same inbred line rescued by either fighter or scrambler males, each of these introduced mutations had to reach a frequency of 5% for them to be retained.

### Statistical analysis

All statistical analysis was performed using R statistical software v.4.4.2 (*84*). Data visualisation carried out using ggplot2 (*85*), mixed effects random models were built using glmmTMB (*86*) and stepwise deletions of non-significant terms performed to reach minimal adequate models. In all missed models block, as described above, was used as random effect to account for the non-independence of multiple GR attempts being performed on the same inbred line and repeated measures of each population.

Generalised linear mixed effects models (GLMMs) were used to analyse total fecundity data (the sum of eggs and larvae in both egg laying vials). These data showed an excess of zeros and therefore analysed using a Tweedie distribution to model both the zeros and non-zeros simultaneously. The full model contained the explanatory variables of GR treatment (no GR, GR from fighters and GR from scramblers) and generation of fecundity assay as a factor (F_1_, F_2_, F_6_, & F_10_), including their interaction term. Analysis was also performed using generation as a continuous variable which showed quantitatively similar results.

The productivity of populations from data collected at F_11_ to F_20_ were analysed using GLMMs with negative binomial error structure. First, to test initial resilience to thermal stress we compared total productivity of each population between the generations when temperature was increased (e.g. F_11_ and F_12_ only), the full model included temperature (which spans across two generations) and GR treatment, including their interaction as explanatory variables. Next, we analysed population productivity of lines only under increased temperature (e.g. F_12_ to F_20_) fitting generation as a continuous variable and GR treatment, including their interaction term as explanatory variables. Population extinction rates were analysed using a mixed effects cox model fitted using coxme (*87*) with GR treatment as an explanatory variable.

We estimated changes in genome-wide genetic variation (π or ϴ) due to GR, Δπ (Δϴ) was calculated as the ratio of π (ϴ) populations with GR at each generation post GR (F_2_, F_6_ and F_10_) divided by the corresponding π (ϴ) of inbred lines at F_0_. Thus, values of 1 would indicate no change in genetic variation, values greater than 1 that genetic variation increased and values less than 1 the genetic variation decreased after GR. GLMMs were then fitted to values of Δπ (Δϴ) with the explanatory variables of generation and GR treatment, including their interaction term.

As described above, fixed deleterious mutations (missense and nonsense) fixed in inbred lines and introduced via GR were determined across entire genomes. We then tested for associations between Δπ and number of each type of mutation separately and whether GR treatment influenced these patterns by fitting GLMMs with Δπ and GR treatment, including their interaction as explanatory variables. We performed the same models but this time using the sum of frequencies of all introduced deleterious mutations as the response variable.

We also determined the amount of fixed deleterious mutations (missense + nonsense) fixed in inbred lines across each chromosome and standardised the amount by chromosome lengths. We then fitted a GLMM to data from all replicates including Δπ of each chromosome and GR treatment, including their interaction term. We further explored these relationships within each individual GR population by performing correlation tests between standardised deleterious load and Δπ on each chromosome. Next, across the entire dataset we compared the mean frequency of initially fixed deleterious (missense or nonsense) at F_10_ that were found on the sex chromosome or autosomes using a GLMM chromosome type as an explanatory variable. To test whether selection was effective at the genome-wide level against initially fixed load we also took the mean frequency of either missense or nonsense mutations at F_10_ and compared these between GR treatments by including it as an explanatory variable in a GLMM. We then calculated the mean frequency of introduced missense or nonsense mutations and compared their relative increase to the relative decrease in mean frequency of initially fixed missense or nonsense mutations (e.g. 1 -mean frequency of introduced mutations) by GLMM, by including GR treatment and introduced/initially fixed mutation type, and their interaction as explanatory variables. To calculate the change in total load in populations we calculated the sum of frequencies of both initially fixed and introduced deleterious mutations (missense or nonsense) at F_10_ and tested whether GR treatment influenced changes in total segregating load by GLMM.

Finally, we combined phenotypic measures and genomic measures to test if genome-wide variation could predict population fitness. We tested if mean inbred line fecundity was predicted by genome-wide π by general linear model. Using GLMMs we tested whether Δπ could explain Δfecundity (mean GR population fecundity / mean inbred line fecundity across all generations) at F_10_ by fitting models with GR treatment and Δπ, including their interaction term as explanatory variables. We also fit a further set of GLMMs including the mean frequency of initially fixed mutations at F_10_ to test whether masking deleterious load could explain any more variance compared to Δπ alone. Further, to explore if inbred line qualities could explain variance in Δfecundity at F_10_, we fit a GLMM that included inbred line genome-wide π, GR treatment and the proportion of fighters in each inbred line replicate at the time of the introduction of rescuer males, including their three-way interaction term. We then tested whether absolute π at F_10_ of GR populations was associated with the generation of extinction (excluding those survived past F_20_) by GLMM, by including GR treatment, genome-wide π and their interaction as explanatory variables. Each population was then assigned a binary factor of either going extinct before or surviving until F_20_ and used GR treatment and absolute π at F_10_, including their interaction term as explanatory variables in a GLMM with binomial error structure.

## Acknowledgements

We would like to thank Sebastian Chmielewski for performing DNA extractions and Agnieszka Szubert-Kruszyńska for assistance in performing experiments.

## Funding

This study was supported by funding from The National Science Centre of Poland 2020/39/D/NZ8/00069 (J.M.P).

## Author contributions

Conceptualization: J.M.P.; Methodology: J.M.P., M.K., J.R.; Investigation: J.M.P, M.K.; Visualization: J.M.P.; Funding acquisition: J.M.P.; Writing – original draft: J.M.P.; Writing – review & editing: J.M.P., M.K., J.R.

## Competing interests

Authors declare that they have no competing interests.

## Appendix

**Fig. S1.**
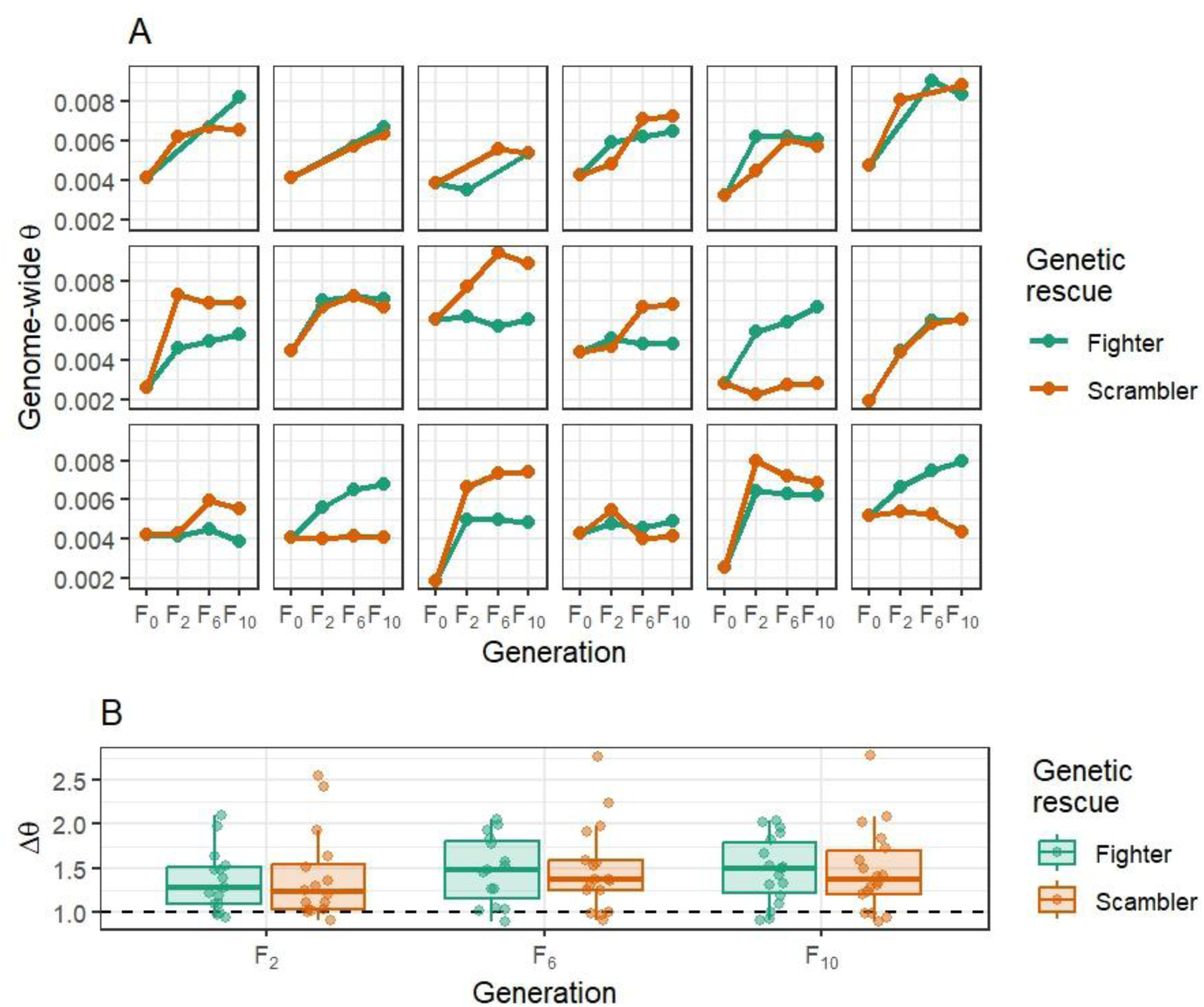
Genome-wide changes in the number of segregating sites (ϴ) due to GR in F_2_, F_6_ and F_10_. **(A)** shows mean genome-wide estimated ϴ from 10kb sliding windows (with 10kb steps) in inbred lines (F_0_) and subsequent generations post GR (F_2_, F_6_ & F_10_). **(B)** represents estimated changes in genome-wide ϴ at each generation post GR after controlling for inbred line measurements (Δ ϴ = ϴ at F_2_ (F_6_ & F_10_) / inbred line ϴ). Thus, a value of 1 for Δϴ (dashed horizonal line) would indicate no change in genome-wide variation, a value less than 1 indicating loss of genetic variation and a value greater than 1 indicating an increase. In both A and B GR populations with rescue from fighters and scramblers are shown in green and orange, respectively. In B boxes are composed of the median and hinge values (25th and 75th percentiles), with whiskers ± interquartile range × 1.5. Individual data points from each population denoted by circles.

**Fig. S2.**
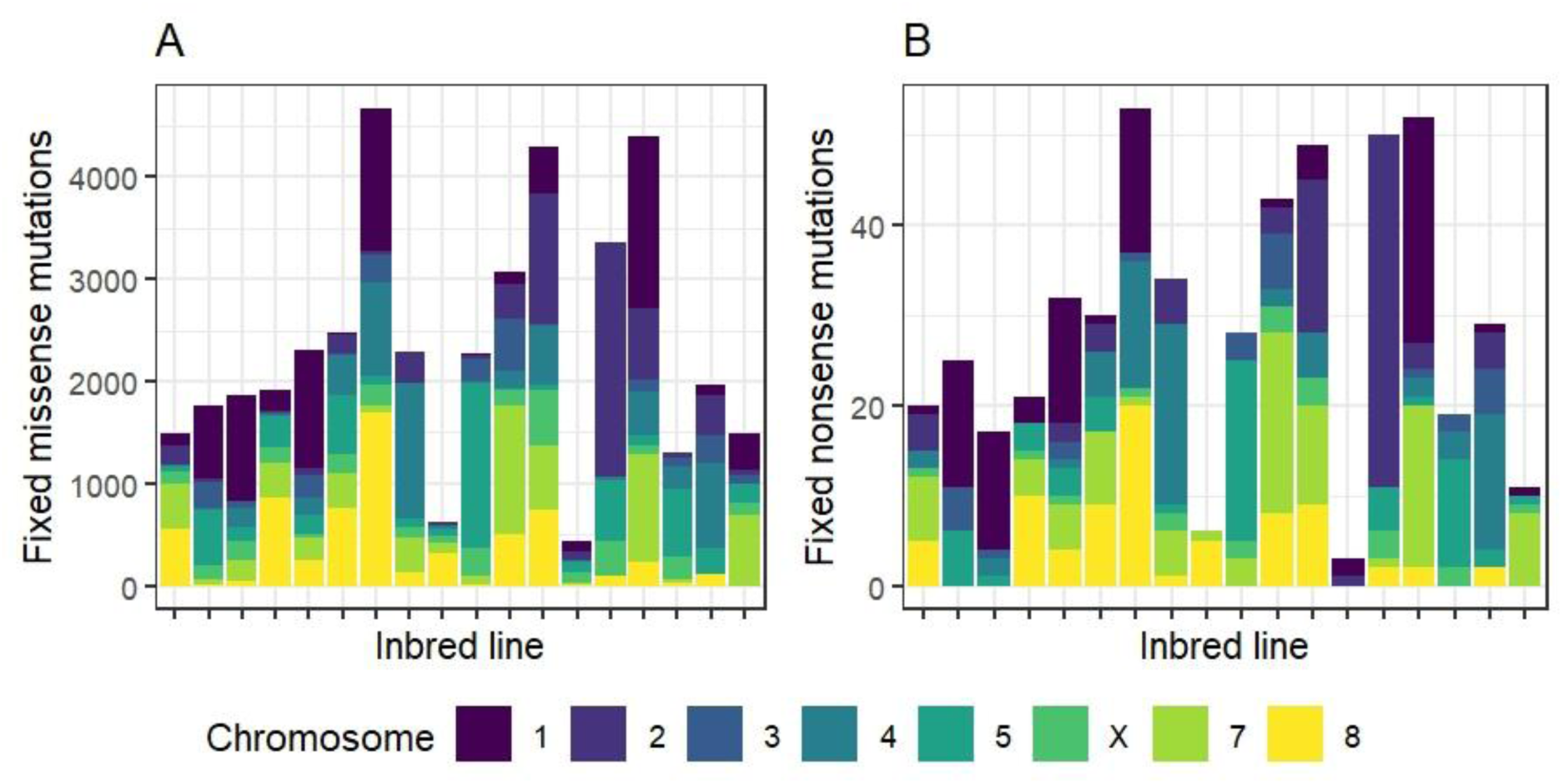
Amount of deleterious load fixed in inbred lines. **(A)** Counts of fixed missense and **(B)** nonsense mutations fixed in each inbred line across each of the chromosomes shown in different colours. In all inbred lines the amount of fixed deleterious mutations (missense and nonsense) was non-randomly distributed across chromosomes (standardised by lengths) by Chi-square test (*p* < 0.001).

**Fig. S3.**
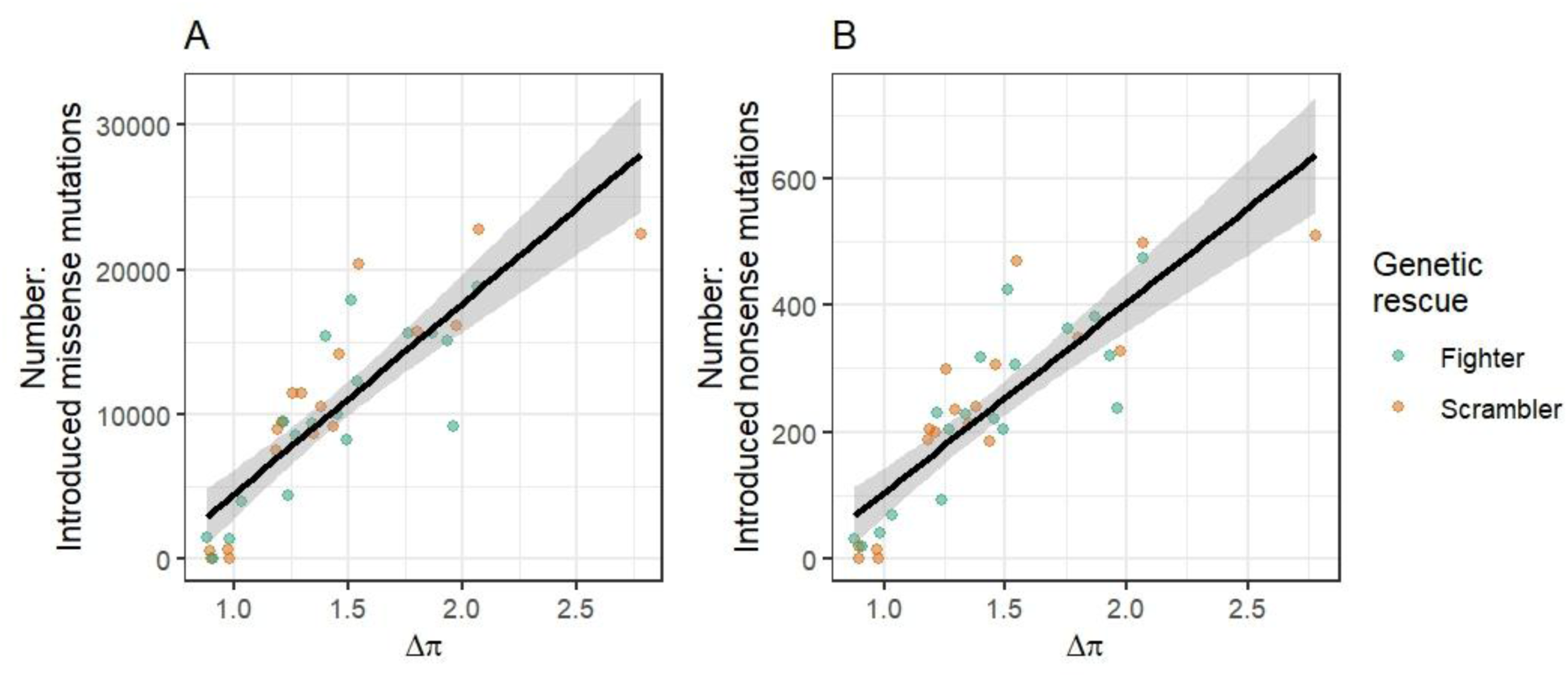
Associations between changes in nucleotide diversity (at F_10_) and number of introduced deleterious mutations. **(A)** number of introduced missense mutations at F_10_. **(B)** number of introduced nonsense mutations at F_10_. In both A and B points are coloured for clarity by whether genetic rescue came from fighter (green) or scrambler (orange) males although in both cases there was no significant effect of male morph on these relationships. Lines indicate line of best of fit and shaded areas the standard error.

**Fig. S4.**
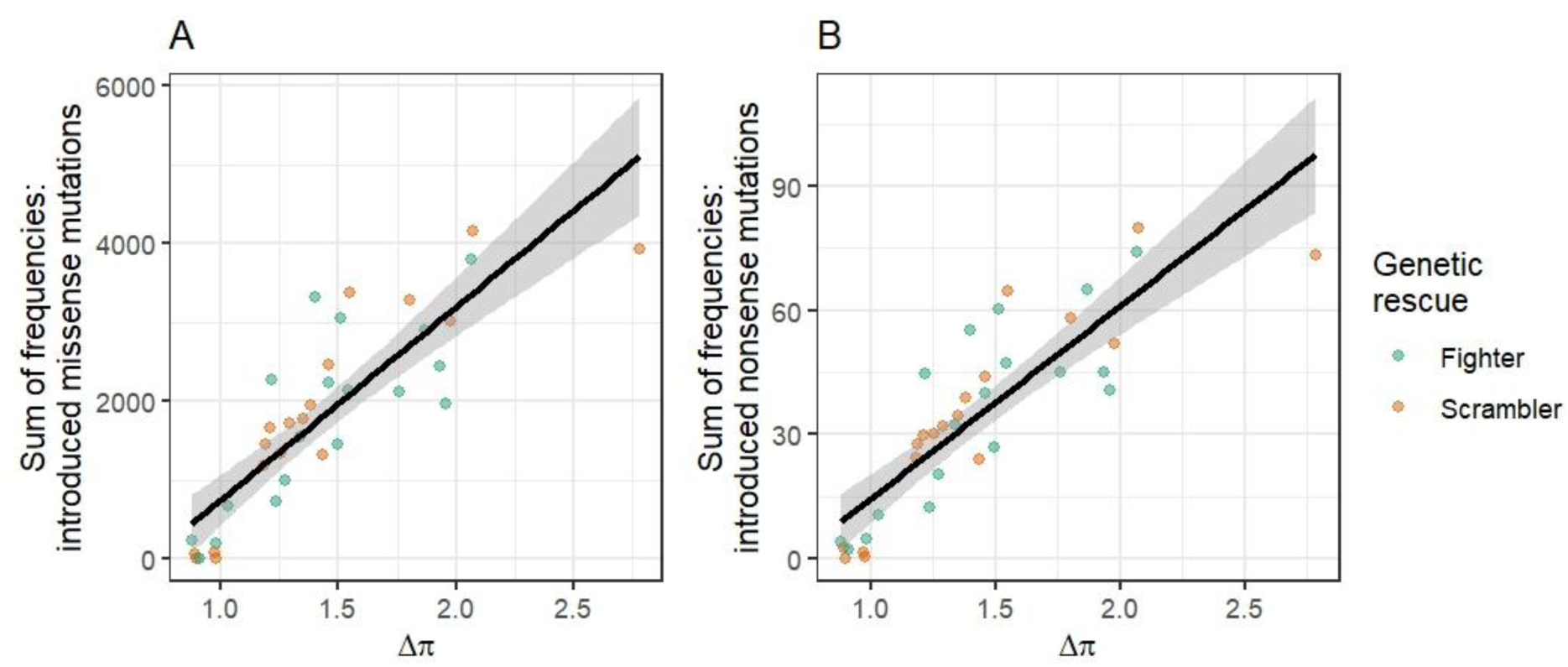
Associations between changes in nucleotide diversity (at F_10_) and the sum of frequencies of introduced deleterious mutations. **(A)** sum of frequencies of introduced missense mutations at F_10_. **(B)** sum of frequencies of introduced nonsense mutations at F_10_. In both A and B points are coloured for clarity by whether genetic rescue came from fighter (green) or scrambler (orange) males although in both cases there was no significant effect of male morph on these relationships. Lines indicate line of best of fit and shaded areas the standard error.

**Fig. S5.**
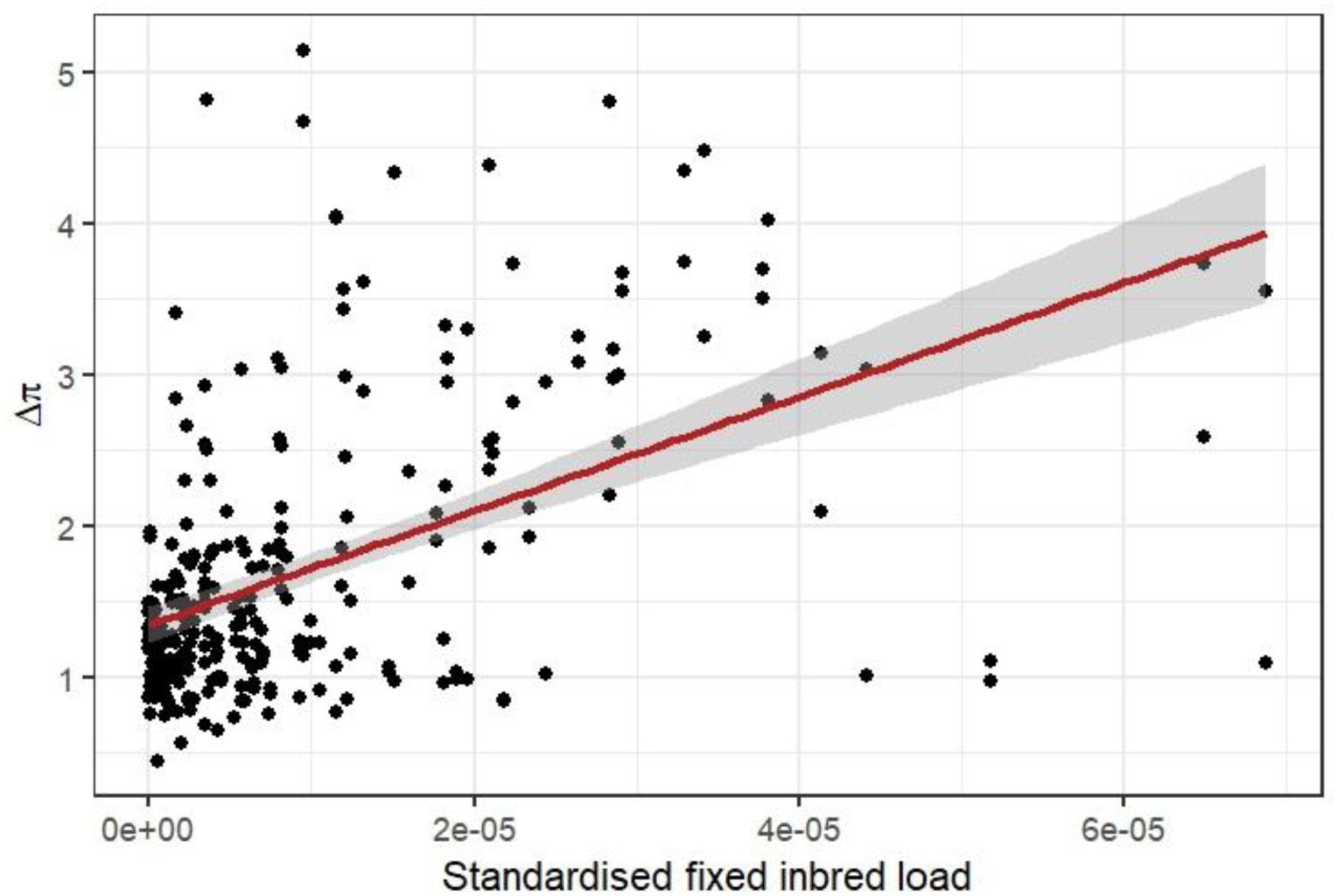
Correlation between fixed load (missense + nonsense) and changes in nucleotide diversity (Δπ) at F_10_ due to GR across all chromosome in the experiment. Association between the number of fixed mutations (both missense and nonsense) on each chromosome (standardised by total chromosome length) from all replicates and the change in nucleotide diversity (Δπ) of that chromosome after GR in F_10_.

**Fig. S6.**
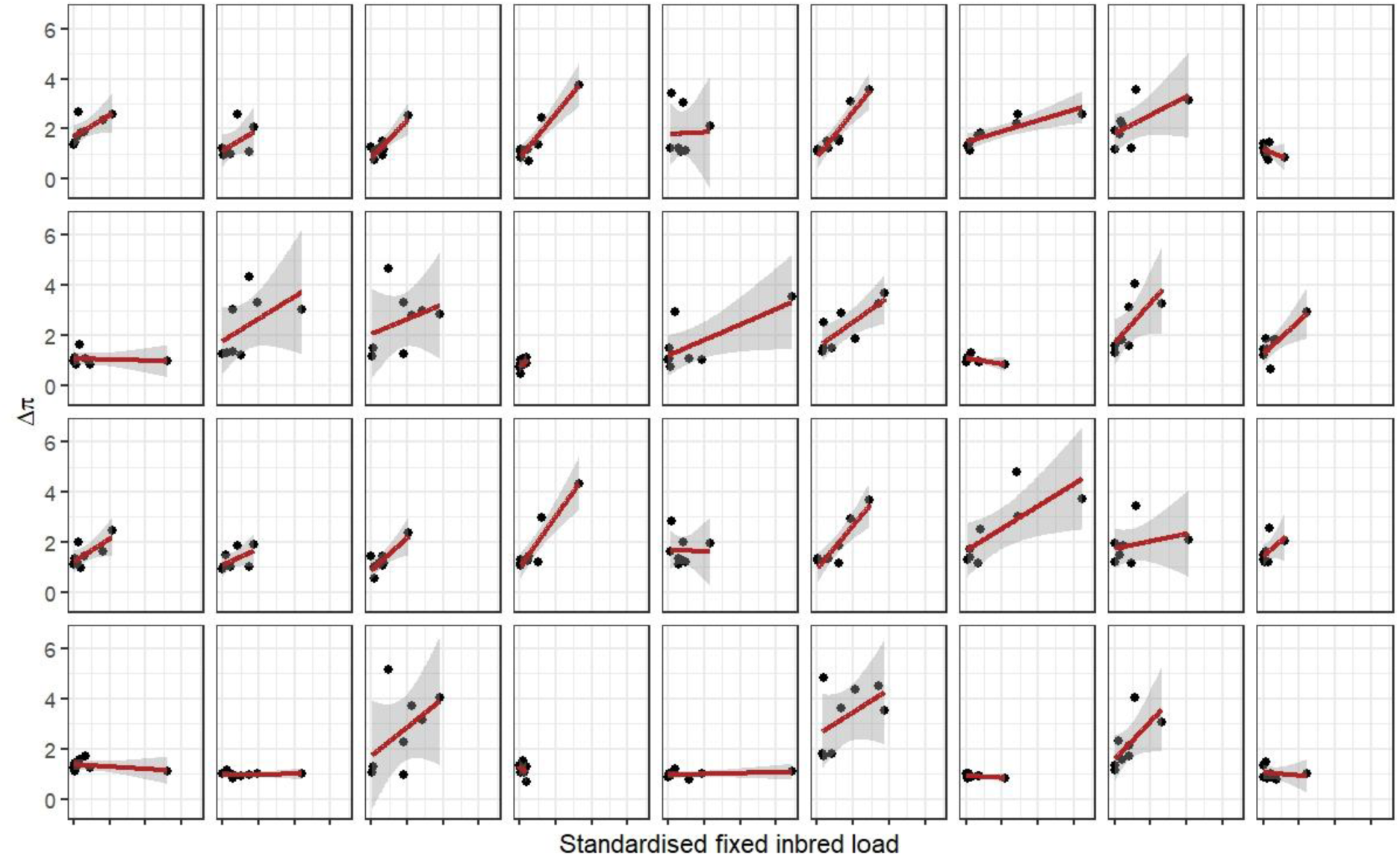
Correlations between fixed load (missense + nonsense) and changes in nucleotide diversity (Δπ) at F_10_ due to GR on each chromosome per replicate. Each panel shows associations between the number of fixed mutations (both missense and nonsense) on each chromosome (standardised by total chromosome length) for each replicate. The two top rows are replicates with GR from fighter males and the two bottom rows are replicates with GR from scrambler males. Lines show fitted line and shaded area the 95% confidence intervals.

**Fig. S7.**
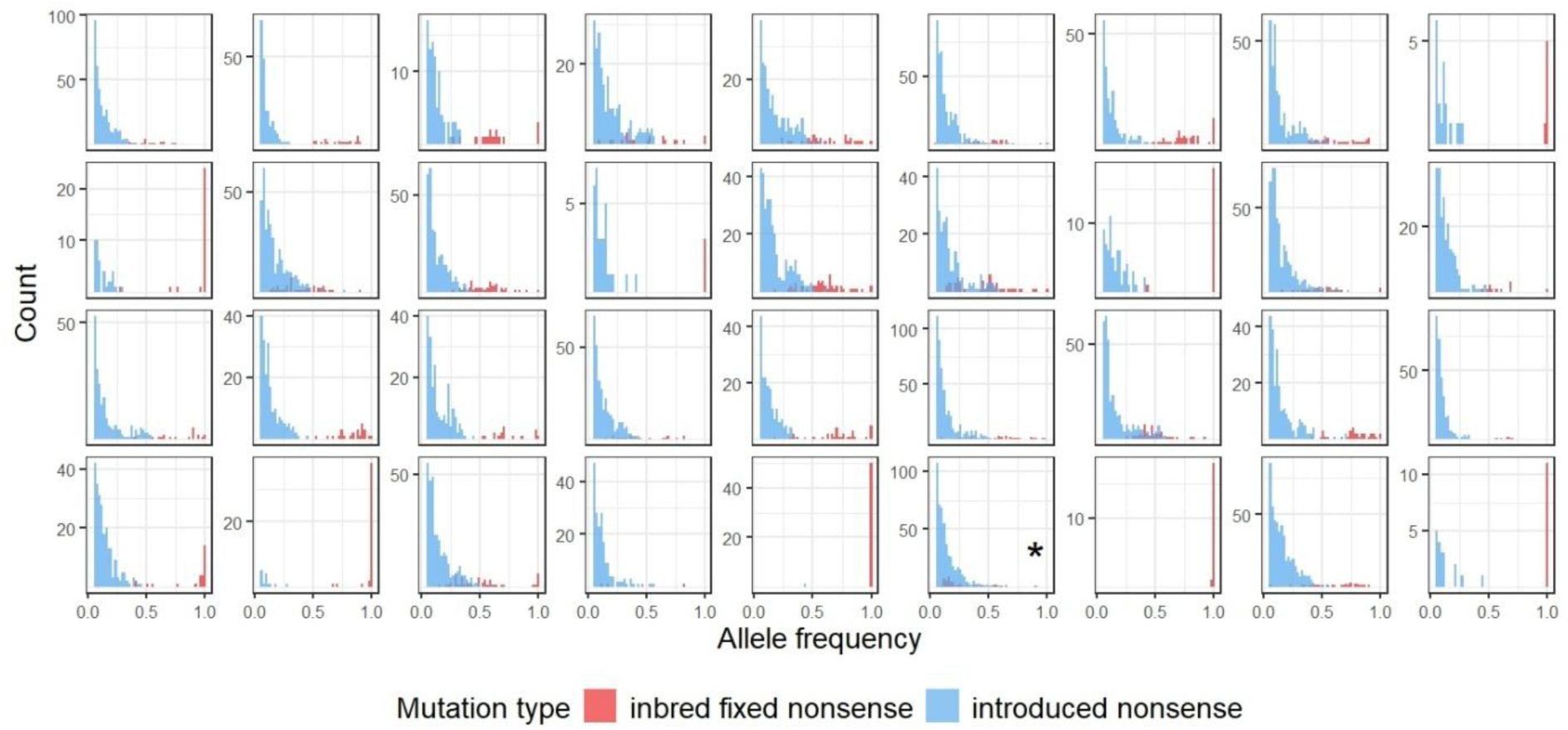
Histograms of allele frequencies of initially fixed and introduced nonsense mutations. Summary histograms of all replicates with GR showing distribution of presumed deleterious nonsense mutations initially fixed in inbred lines (red) and introduced via GR (blue) at F_10_ from across all eight chromosomes. Histograms in the two top rows are replicates with GR from fighter males and the two bottom rows are replicates with GR from scrambler males. The plot highlighted with an asterisk (*) is the example replicate from Fig. 2a in main text.

**Fig. S8.**
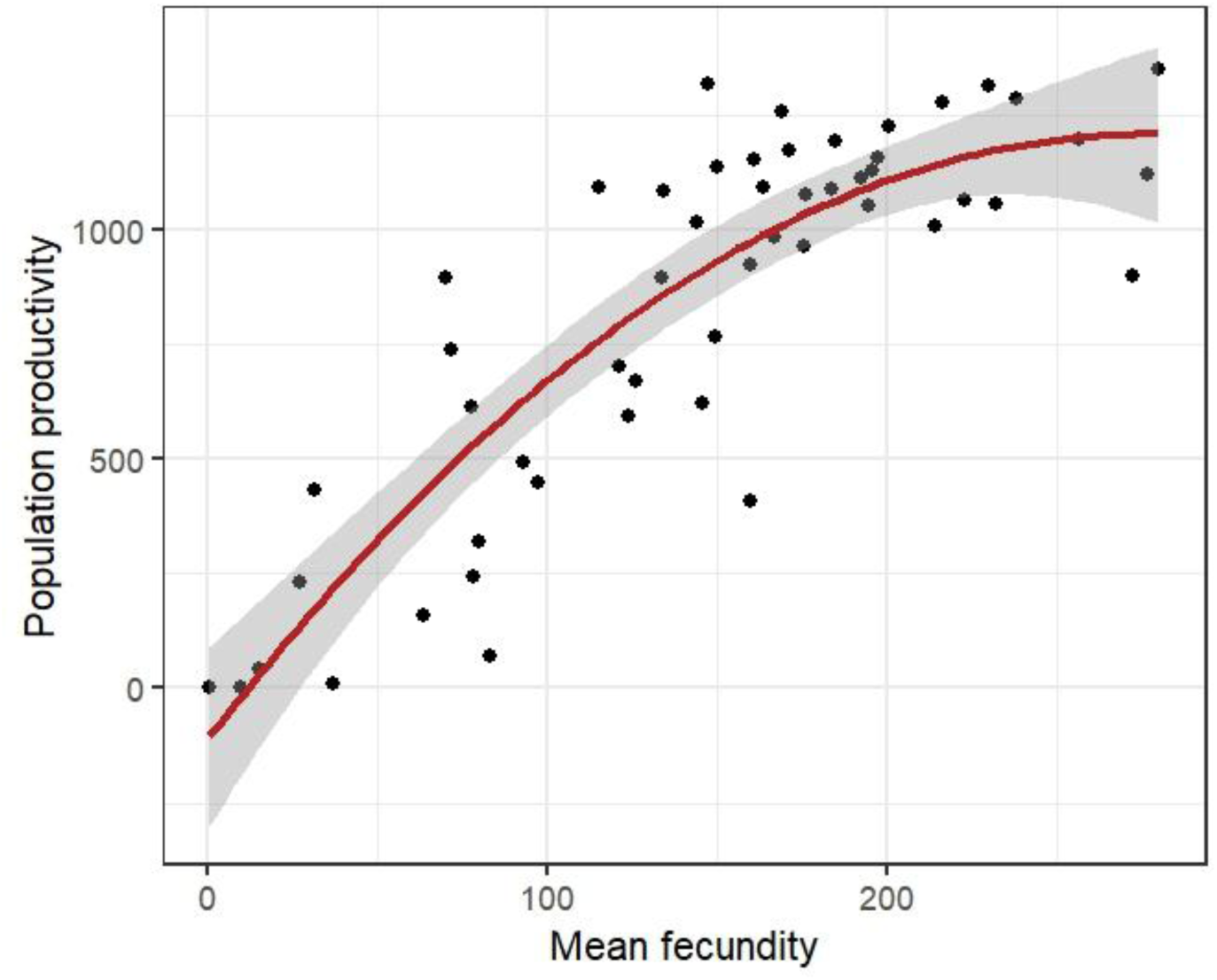
Fecundity of male and female pairs predicts productivity of populations. The association between mean fecundity of male and female pairs in each replicate at F_10_ with the total number of adults and tritonymphs produced by populations at F_11_. The quadratic term was significant (*χ^2^* = 32.85, *df* = 1, *p* < 0.001) indicating that the relationship is non-linear and likely a consequence of density effects during rearing. The line represents line of best fit (including the quadratic term) and shaded area the standard error.

**Fig. S9.**
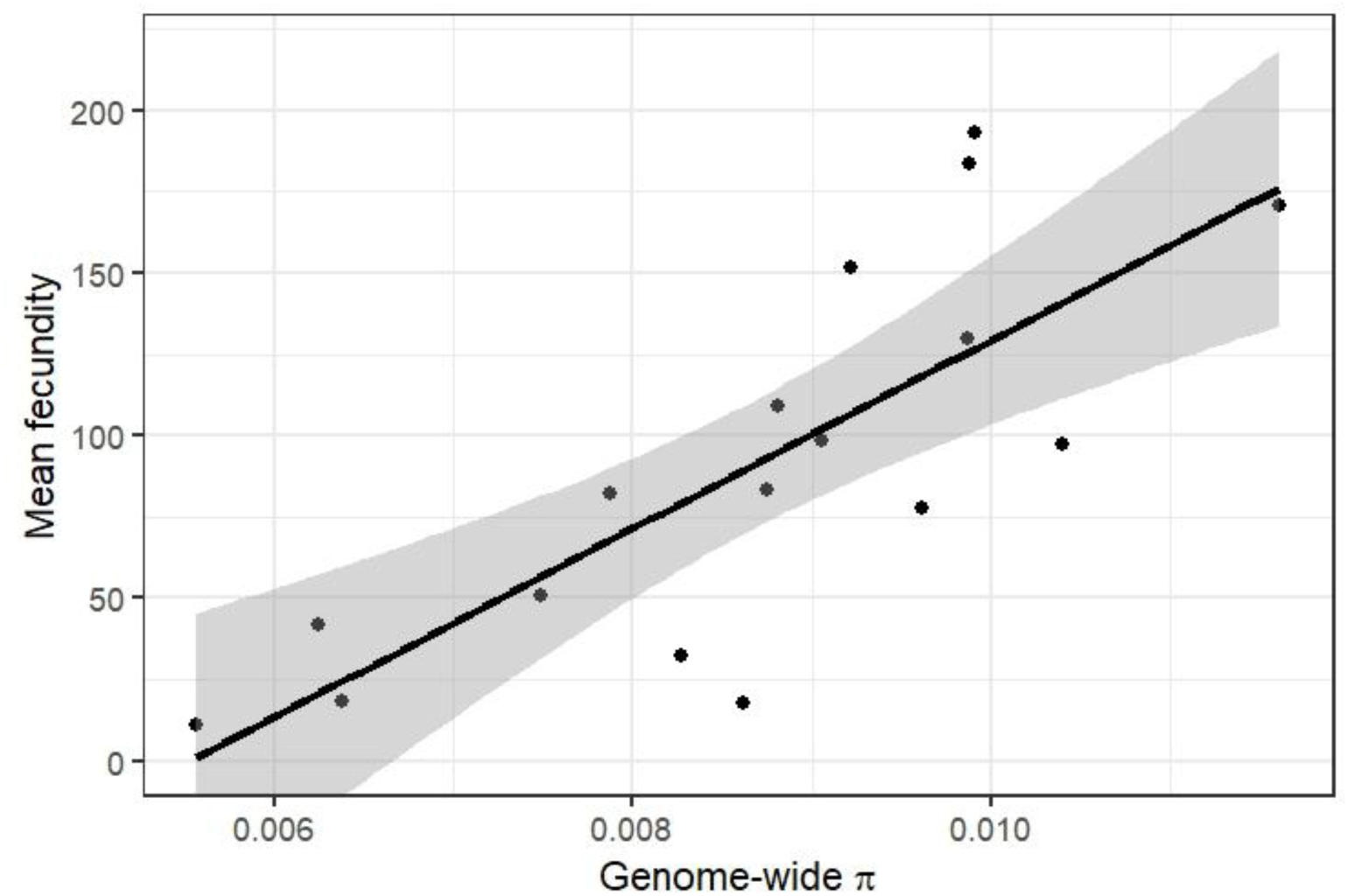
Association between genome-wide nucleotide diversity (π) of inbred lines and fecundity. Each point represents the mean fecundity of male and female across all generations from inbred lines and their corresponding genome-wide π. There was a significant positive association between genome-wide π and mean inbred line fecundity (F = 23.26, df = 1, *p* < 0.001). The line represents line of best fit and shaded areas the standard error.

**Fig. S10.**
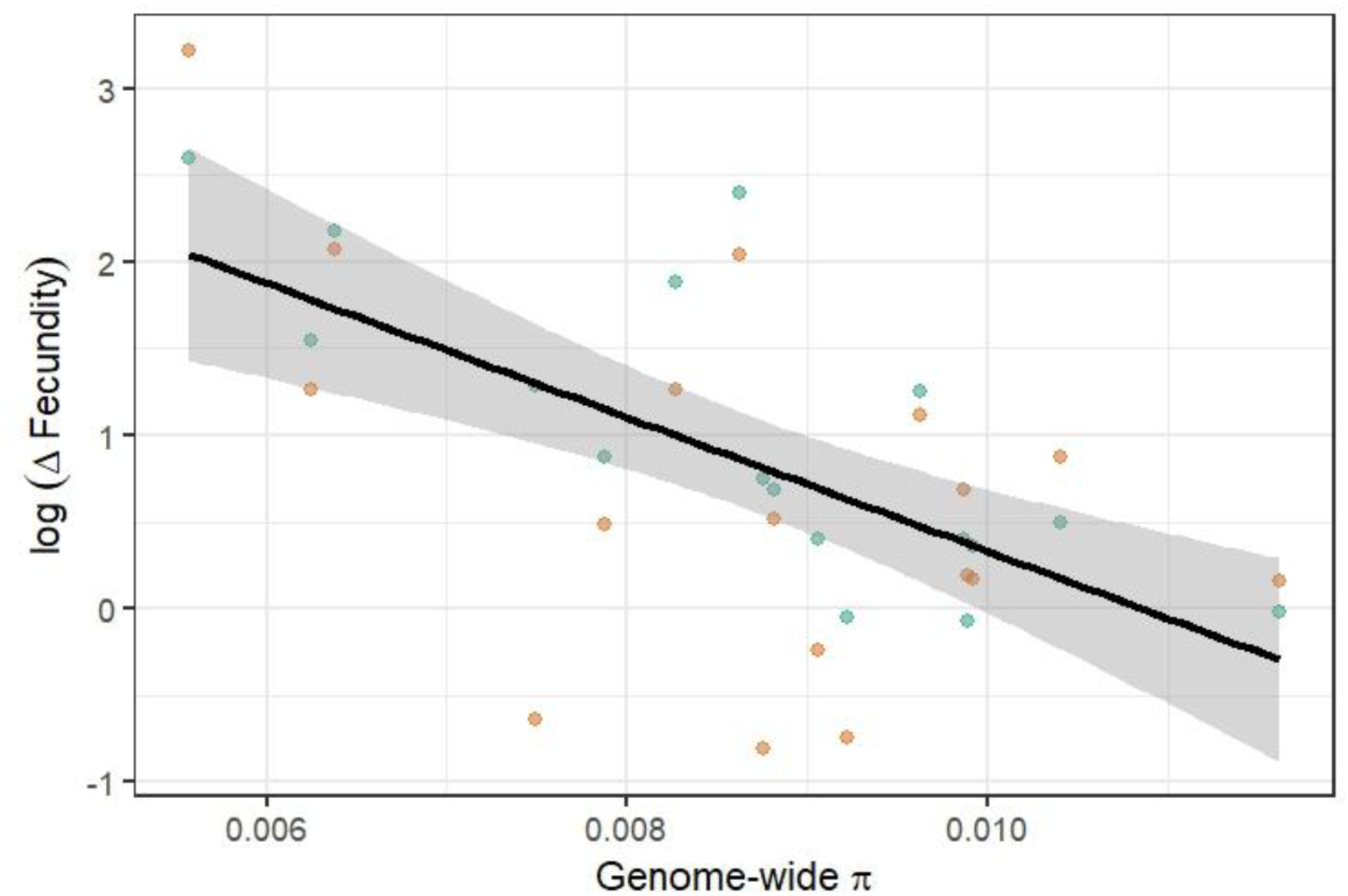
Associations between genome-wide nucleotide diversity (π) of inbred lines and relative change in fecundity due to GR at F_10_. Each point represents the relative change in mean fecundity of male and female at F_10_ (Δfecundity = mean fecundity of GR populations at F_10_ / mean fecundity of inbred lines across all generations) and their corresponding genome-wide π of inbred lines at F_0_. Points are coloured for clarity by whether genetic rescue came from fighter (green) or scrambler (orange) males although there was no significant effect of male morph on this relationship. The line indicates line of best of fit and shaded area the standard error.

**Fig. S11.**
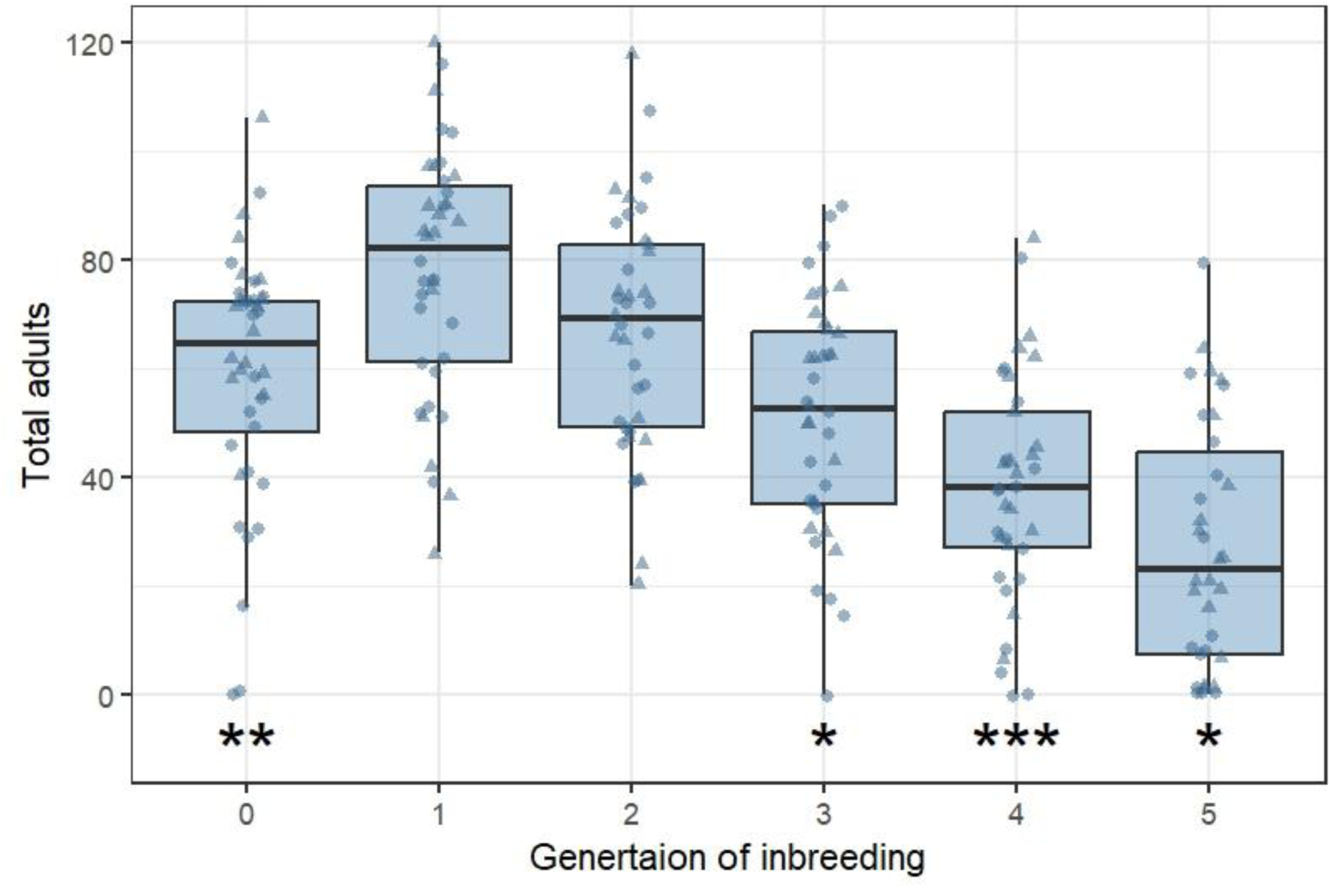
Decline in population fitness as inbreeding increases. Total number of surviving adults each generation after full sib mating plotting only pairs which were used in following generations. Boxes are composed of the median and hinge values (25th and 75th percentiles), with whiskers ± interquartile range × 1.5. Individual replicates are denoted by circles and triangles, the latter being those inbred lines that were used in the full GR experiment. The asterisk at bottom of plots indicate the number of inbred lines that went extinct at that generation.

**Table S1.** Model selection process and summary of minimal adequate model of changes in π (Δπ) across the three generations of genomic sampling (F_2_, F_6_ & F_10_). GLMMs were fit with a Gaussian error structure.

| Model selection |  |  |  |  |
| --- | --- | --- | --- | --- |
| Source of variation | $\chi^2$ | d.f. | p-value | |
| GR | 0.232 | 1 | 0.630 |  |
| Generation | 11.403 | 2 | 0.003 |  |
| GR : Generation | 0.677 | 2 | 0.713 |  |
| Minimal adequate model |  |  |  |  |
| Random effects | Variance |  | Standard deviation |  |
| Block (e.g. inbred line) | 0.126 |  | 0.355 |  |
|  | Observations: n = 99 |  | Blocks: n = 18 |  |
| Explanatory variable | Estimate | S.E. | z-value | p-value |
| Intercept | 1.272 | 0.091 | 13.91 | < 0.001 |
| Generation F<sub>6</sub> | 0.147 | 0.051 | 2.90 | 0.004 |
| Generation F<sub>10</sub> | 0.159 | 0.050 | 3.19 | 0.001 |

d.f. = degrees of freedom; S.E. = standard error; GR = genetic rescue treatment
Intercept contains GR – fighter and Generation F<sub>2</sub>

**Table S2.**
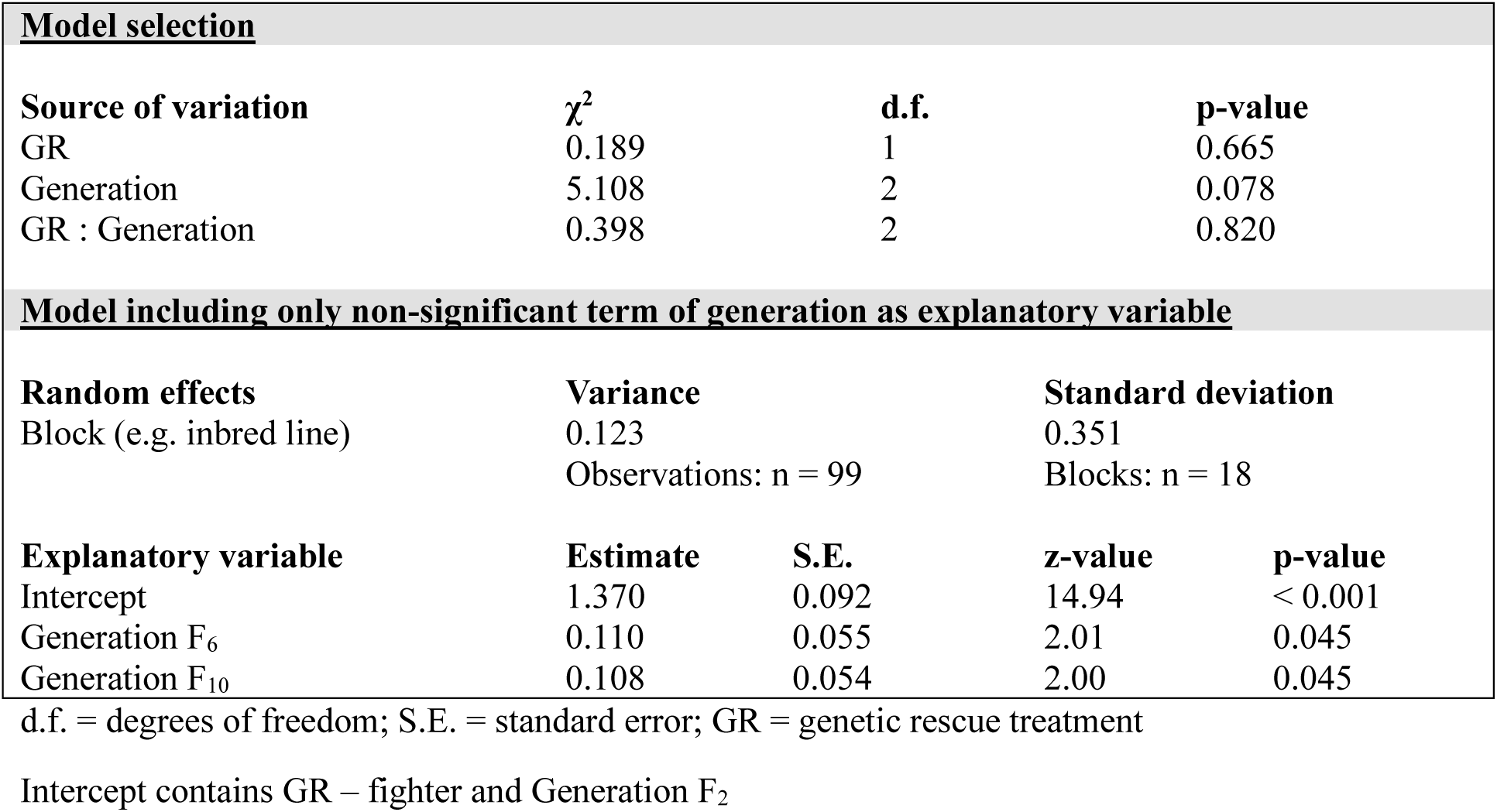
Model selection process and model summary changes in π (Δπ) across the three generations of genomic sampling (F_2_, F_6_ & F_10_). GLMMs were fit with a Gaussian error structure. Note that the effect of generation was non-significant (*p* = 0.078) but included for comparison with GLMM of Δπ (Table S1).

**Table S3.** Model selection process and summaries of minimal adequate models of number of introduced missense or nonsense mutations at F_10_. GLMMs were fit with negative binomial error structure.

| Model selection: missense |  |  |  |  |
| --- | --- | --- | --- | --- |
| Source of variation | $\chi^2$ | d.f. | p-value | |
| GR | 0.042 | 1 | 0.838 |  |
| $\Delta\pi$ | 13.621 | 1 | < **0.001** | |
| GR : $\Delta\pi$ | 0.060 | 1 | 0.806 | |
| Minimal adequate model: missense |  |  |  |  |
| Random effects | Variance |  | Standard deviation |  |
| Block (e.g. inbred line) | 2.18e-09 |  | 4.669e-05 |  |
|  | Observations: n = 36 |  | Blocks: n = 18 |  |
| Explanatory variable | Estimate | S.E. | z-value | p-value |
| Intercept | 6.351 | 0.780 | 8.14 | < **0.001** |
| $\Delta\pi$ | 1.882 | 0.534 | 5.52 | < **0.001** |
| Model selection: nonsense |  |  |  |  |
| Source of variation | $\chi^2$ | d.f. | p-value | |
| GR | 0.014 | 1 | 0.905 |  |
| $\Delta\pi$ | 16.750 | 1 | < **0.001** | |
| GR : $\Delta\pi$ | 1.152 | 1 | 0.697 | |
| Minimal adequate model: nonsense |  |  |  |  |
| Random effects | Variance |  | Standard deviation |  |
| Block (e.g. inbred line) | 1.194e-09 |  | 3.456e-05 |  |
|  | Observations: n = 36 |  | Blocks: n = 18 |  |
| Explanatory variable | Estimate | S.E. | z-value | p-value |
| Intercept | 2.678 | 0.650 | 4.12 | < **0.001** |
| $\Delta\pi$ | 1.816 | 0.445 | 4.09 | < **0.001** |

d.f. = degrees of freedom; S.E. = standard error; GR = genetic rescue treatment; $\Delta\pi$ = change in $\pi$ between inbred line and F<sub>10</sub>

**Table S4.** – Model selection process and summaries of minimal adequate models of the sum of frequencies of introduced missense or nonsense mutations at F_10_. GLMMs were fit with negative binomial error structure.

| Model selection: missense |  |  |  |  |
| --- | --- | --- | --- | --- |
| Source of variation | $\chi^2$ | d.f. | p-value | |
| GR | 0.093 | 1 | 0.760 |  |
| $\Delta\pi$ | 45.936 | 1 | < 0.001 | |
| GR : $\Delta\pi$ | 0.015 | 1 | 0.901 | |
| Minimal adequate model: missense |  |  |  |  |
| Random effects | Variance |  | Standard deviation |  |
| Block (e.g. inbred line) | 1.811e-29 |  | 4.255e-15 |  |
|  | Observations: n = 36 |  | Blocks: n = 18 |  |
| Explanatory variable | Estimate | S.E. | z-value | p-value |
| Intercept | -1705.4 | 378.7 | -4.50 | < 0.001 |
| $\Delta\pi$ | 2450.4 | 254.1 | 9.64 | < 0.001 |
| Model selection: nonsense |  |  |  |  |
| Source of variation | $\chi^2$ | d.f. | p-value | |
| GR | 0.002 | 1 | 0.957 |  |
| $\Delta\pi$ | 47.536 | 1 | < 0.001 | |
| GR : $\Delta\pi$ | 0.109 | 1 | 0.741 | |
| Minimal adequate model: nonsense |  |  |  |  |
| Random effects | Variance |  | Standard deviation |  |
| Block (e.g. inbred line) | 52.63 |  | 7.255 |  |
|  | Observations: n = 36 |  | Blocks: n = 18 |  |
| Explanatory variable | Estimate | S.E. | z-value | p-value |
| Intercept | -36.704 | 8.376 | -4.382 | < 0.001 |
| $\Delta\pi$ | 49.849 | 5.624 | 8.863 | < 0.001 |

d.f. = degrees of freedom; S.E. = standard error; GR = genetic rescue treatment; $\Delta\pi$ = change in $\pi$ between inbred line and F<sub>10</sub>

**Table S5.** Model selection process and summary of minimal adequate model of standardised fixed deleterious load on each chromosome. GLMMs were fit with a Gaussian error structure.

| Model selection |  |  |  |  |
| --- | --- | --- | --- | --- |
| Source of variation | $\chi^2$ | d.f. | p-value | |
| GR | 0.351 | 1 | 0.554 |  |
| Std. load | 71.225 | 1 | < 0.001 |  |
| GR : Std. load | 0.418 | 1 | 0.518 |  |
| Model including only non-significant term of generation as explanatory variable |  |  |  |  |
| Random effects | Variance |  | Standard deviation |  |
| Block (e.g. inbred line) | 0.140 |  | 0.375 |  |
|  | Observations: n = 288 |  | Blocks: n = 18 |  |
| Explanatory variable | Estimate | S.E. | z-value | p-value |
| Intercept | 1.393e+00 | 1.032e-01 | 13.50 | < 0.001 |
| Std. load | 3.294e+04 | 3.687e+03 | 8.93 | < 0.001 |

d.f. = degrees of freedom; S.E. = standard error; GR = genetic rescue treatment; Std. load = Number of fixed deleterious mutations (missense + nonsense) standardised by chromosome length

**Table S6.**
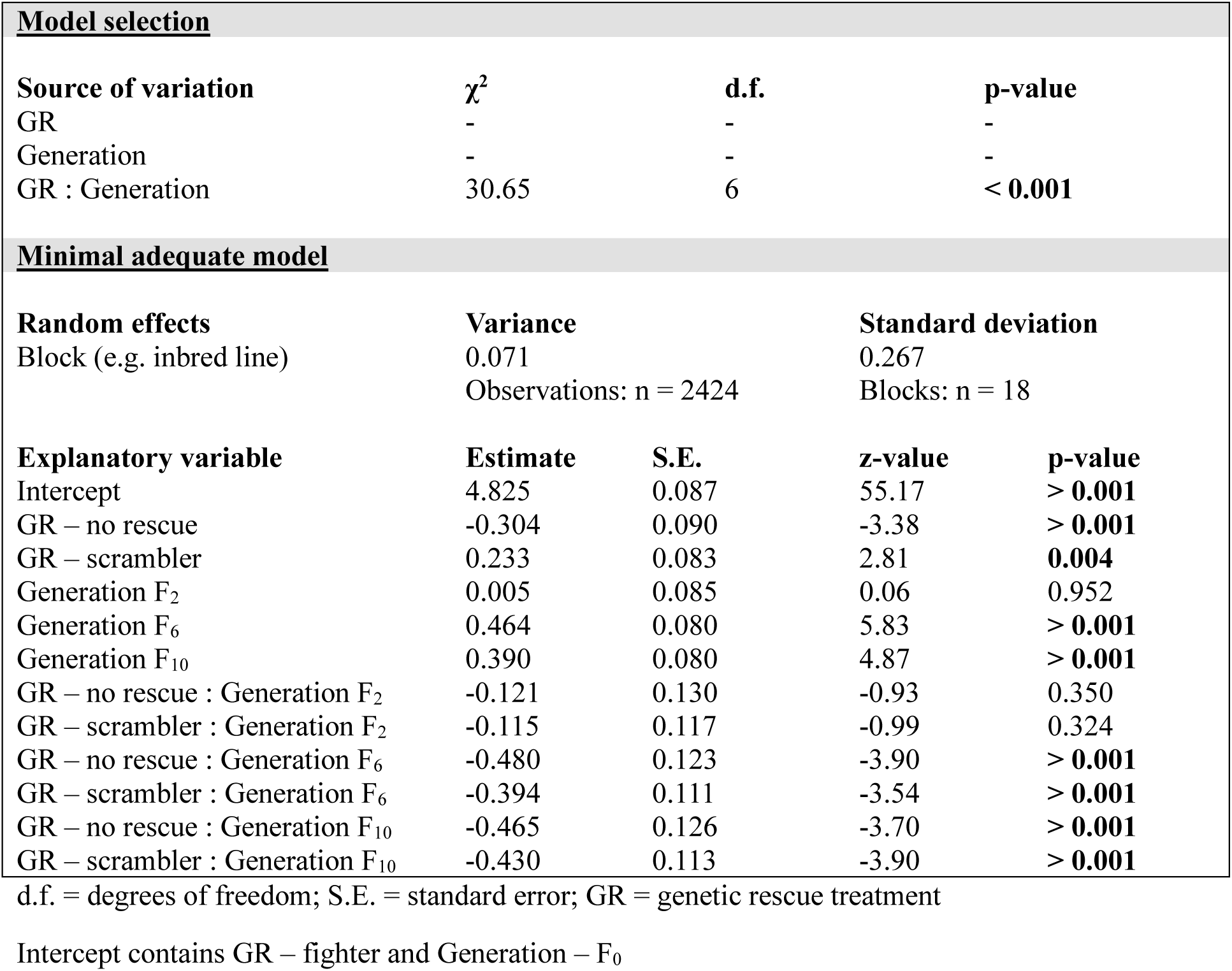
Model selection process and summary of minimal adequate model of fecundity of male and female pairs. Fecundity was determined across nine days of egg laying. GLMMs were fit using a Tweedie error structure to model both non-zeros and the excess of zeros simultaneously.

| Model selection |  |  |  |  |
| --- | --- | --- | --- | --- |
| Source of variation | $\chi^2$ | d.f. | p-value | |
| GR | - | - | - |  |
| Generation | - | - | - |  |
| GR : Generation | 30.65 | 6 | < 0.001 |  |
| Minimal adequate model |  |  |  |  |
| Random effects | Variance | Standard deviation |  |  |
| Block (e.g. inbred line) | 0.071 | 0.267 |  |  |
|  | Observations: n = 2424 | Blocks: n = 18 |  |  |
| Explanatory variable | Estimate | S.E. | z-value | p-value |
| Intercept | 4.825 | 0.087 | 55.17 | > 0.001 |
| GR – no rescue | -0.304 | 0.090 | -3.38 | > 0.001 |
| GR – scrambler | 0.233 | 0.083 | 2.81 | 0.004 |
| Generation F<sub>2</sub> | 0.005 | 0.085 | 0.06 | 0.952 |
| Generation F<sub>6</sub> | 0.464 | 0.080 | 5.83 | > 0.001 |
| Generation F<sub>10</sub> | 0.390 | 0.080 | 4.87 | > 0.001 |
| GR – no rescue : Generation F<sub>2</sub> | -0.121 | 0.130 | -0.93 | 0.350 |
| GR – scrambler : Generation F<sub>2</sub> | -0.115 | 0.117 | -0.99 | 0.324 |
| GR – no rescue : Generation F<sub>6</sub> | -0.480 | 0.123 | -3.90 | > 0.001 |
| GR – scrambler : Generation F<sub>6</sub> | -0.394 | 0.111 | -3.54 | > 0.001 |
| GR – no rescue : Generation F<sub>10</sub> | -0.465 | 0.126 | -3.70 | > 0.001 |
| GR – scrambler : Generation F<sub>10</sub> | -0.430 | 0.113 | -3.90 | > 0.001 |

d.f. = degrees of freedom; S.E. = standard error; GR = genetic rescue treatment
Intercept contains GR – fighter and Generation – F<sub>0</sub>

**Table S7.** Model selection process and summary of minimal adequate model of changes in mean fecundity (Δfecundity) at F_10_. GLMMs were fit using Gaussian error structure.

| Model selection |  |  |  |  |
| --- | --- | --- | --- | --- |
| Source of variation | $\chi^2$ | d.f. | p-value | |
| GR | 14.316 | 1 | < 0.001 |  |
| $\Delta\pi$ | 50.435 | 1 | < 0.001 | |
| GR : $\Delta\pi$ | 1.101 | 1 | 0.294 | |
| Minimal adequate model |  |  |  |  |
| Random effects | Variance |  | Standard deviation |  |
| Block (e.g. inbred line) | 0.248 |  | 0.498 |  |
|  | Observations: n = 34 |  | Blocks: n = 17 |  |
| Explanatory variable | Estimate | S.E. | z-value | p-value |
| Intercept | -1.278 | 0.248 | -5.14 | < 0.001 |
| GR – scrambler | 1.625 | 0.152 | 10.70 | < 0.001 |
| $\Delta\pi$ | -0.295 | 0.061 | -4.83 | < 0.001 |

d.f. = degrees of freedom; S.E. = standard error; GR = genetic rescue treatment; $\Delta\pi$ = change in $\pi$ between inbred line and F<sub>10</sub>
Intercept contains GR – fighter

**Table S8.** Model selection process and summary of minimal adequate model of changes in mean fecundity (Δfecundity) at F_10_. GLMMs were fit using Gaussian error structure.

| <u>Model selection</u> |  |  |  |  |
| --- | --- | --- | --- | --- |
| Source of variation | $\chi^2$ | d.f. | p-value | |
| GR | 0.097 | 1 | 0.755 |  |
| Inbred $\pi$ | 10.203 | 1 | **0.001** | |
| Resident MR | 1.670 | 1 | 0.196 |  |
| GR : Inbred $\pi$ | 0.026 | 1 | 0.871 | |
| GR : Resident MR | 0.114 | 1 | 0.915 |  |
| Inbred $\pi$ : Resident MR | 0.000 | 1 | 0.984 | |
| GR : Inbred $\pi$ : Resident MR | 2.916 | 1 | 0.088 | |
| <u>Minimal adequate model</u> |  |  |  |  |
| Random effects | Variance |  | Standard deviation |  |
| Block (e.g. inbred line) | 0.315 |  | 0.561 |  |
|  | Observations: n = 136 |  | Blocks: n = 17 |  |
| Explanatory variable | Estimate | S.E. | z-value | p-value |
| Intercept | 3.732 | 0.806 | 4.63 | < **0.001** |
| Inbred $\pi$ | -341.649 | 91.368 | -3.74 | < **0.001** |

d.f. = degrees of freedom; S.E. = standard error; GR = genetic rescue treatment; Inbred $\pi$ = Nucleotide diversity of inbred line; Resident MR = Morph proportion in inbred line at time of introduction

**Table S9.**
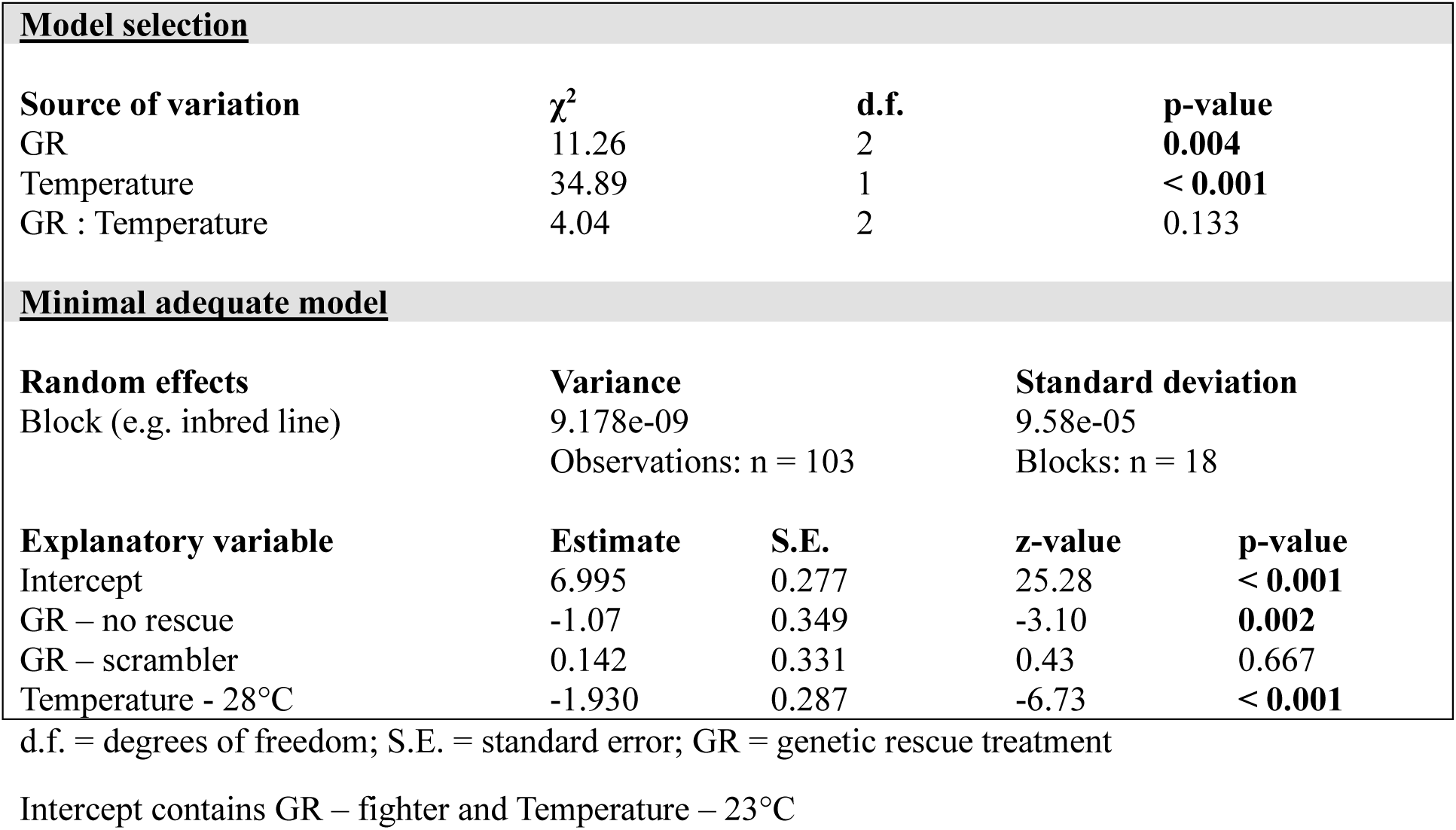
Model selection process and summary of minimal adequate model of population productivity as temperature was increased by 5°C. Note that the temperature explanatory variable is also associated with generation of experiment (e.g. 23°C at F_11_ and 28°C at F_12_). GLMMs were fit using negative binomial error structure.

**Table S10.** Model selection process and summary of minimal adequate model of population productivity during the extinction assay. Generation was fit as continuous variable from F_12_ to F_20_ and GLMMs were fit using negative binomial error structure.

| <u>Model selection</u> |  |  |  |  |
| --- | --- | --- | --- | --- |
| Source of variation | $\chi^2$ | d.f. | p-value | |
| GR | 22.69 | 2 | < **0.001** |  |
| Generation | 6.30 | 1 | **0.012** |  |
| GR : Generation | 3.69 | 2 | 0.158 |  |
| <u>Minimal adequate model</u> |  |  |  |  |
| Random effects | Variance |  | Standard deviation |  |
| Block (e.g. inbred line) | 0.228 |  | 0.477 |  |
|  | Observations: n = 240 |  | Blocks: n = 18 |  |
| Explanatory variable | Estimate | S.E. | z-value | p-value |
| Intercept | 6.074 | 0.688 | 8.83 | < **0.001** |
| GR – no rescue | -1.930 | 0.436 | -4.43 | < **0.001** |
| GR – scrambler | 0.288 | 0.270 | 1.06 | 0.286 |
| Generation | -0.113 | 0.044 | -2.58 | **0.010** |

d.f. = degrees of freedom; S.E. = standard error; GR = genetic rescue treatment
Intercept contains GR – fighter and Generation – F<sub>12</sub>

**Table S11.** Summary of Cox proportional hazard model assessing extinctions of populations across the entire experiment. Model was fit using a mixed effects Cox model.

| **Model summary** |  |  |  |  |
| --- | --- | --- | --- | --- |
| **Random effects** | **Variance** |  | **Standard deviation** |  |
| Block (e.g. inbred line) | 9.047e-03 |  | 8.186e-05 |  |
|  | Events: n = 43, 54 |  | Blocks: n = 18 |  |
| **Explanatory variable** | **Estimate** | **exp(coef)** | **S.E.(coef)** | **p-value** |
| GR – no rescue | 1.625 | 5.080 | 0.397 | < **0.001** |
| GR – scrambler | -0.093 | 0.911 | 0.402 | 0.820 |

S.E. = standard error; coef = coefficient; GR = genetic rescue treatment
Comparison made against GR – fighter

**Table S12.** Model selection process and summary of minimal adequate model of generation of population extinction. GLMMs were fit using Gaussian error structure.

| Model selection |  |  |  |  |
| --- | --- | --- | --- | --- |
| Source of variation | $\chi^2$ | d.f. | p-value | |
| GR | 0.947 | 1 | 0.330 |  |
| F<sub>10</sub> $\pi$ | 22.278 | 1 | < **0.001** | |
| GR : F<sub>10</sub> $\pi$ | 0.111 | 1 | 0.740 | |
| Minimal adequate model |  |  |  |  |
| Random effects | Variance |  | Standard deviation |  |
| Block (e.g. inbred line) | 0.623 |  | 0.789 |  |
|  | Observations: n = 23 |  | Blocks: n = 15 |  |
| Explanatory variable | Estimate | S.E. | z-value | p-value |
| Intercept | 7.395 | 1.163 | 6.65 | < **0.001** |
| F<sub>10</sub> $\pi$ | 671.102 | 99.153 | 6.768 | < **0.001** |

d.f. = degrees of freedom; S.E. = standard error; GR = genetic rescue treatment; F<sub>10</sub> $\pi$ = absolute nucleotide diversity at F<sub>10</sub>

**Table S13.** Model selection process and summary of minimal adequate model of populations that went extinct before or survived until F_20_. GLMMs were fit using binomial error structure.

| Model selection |  |  |  |  |
| --- | --- | --- | --- | --- |
| Source of variation | $\chi^2$ | d.f. | p-value | |
| GR | 1.178 | 1 | 0.278 |  |
| F<sub>10</sub> $\pi$ | 10.829 | 1 | < **0.001** | |
| GR : F<sub>10</sub> $\pi$ | 0.689 | 1 | 0.407 | |
| Minimal adequate model |  |  |  |  |
| Random effects | Variance |  | Standard deviation |  |
| Block (e.g. inbred line) | 1.253e-49 |  | 3.54e-25 |  |
|  | Observations: n = 36 |  | Blocks: n = 18 |  |
| Explanatory variable | Estimate | S.E. | z-value | p-value |
| Intercept | -9.381 | 3.393 | -2.77 | **0.006** |
| F<sub>10</sub> $\pi$ | 726.720 | 272.103 | 2.671 | **0.008** |

d.f. = degrees of freedom; S.E. = standard error; GR = genetic rescue treatment; F<sub>10</sub> $\pi$ = absolute nucleotide diversity at F<sub>10</sub>

## Notes

### Competing Interest Statement

The authors have declared no competing interest.

## References

1. M. R. W. Rands et al., Science (1979). 329, 1298–1303 (2010).

2. B. W. Brook, N. S. Sodhi, C. J. A. Bradshaw, Trends Ecol Evol. 23, 453–460 (2008).

3. S. H. M. Butchart et al., Science (1979). 328, 1164–1168 (2010).

4. K. R. Crooks et al., PNAS. 114, 7635–7640 (2017).

5. N. M. Haddad et al., Sci Adv. 1, 1–9 (2015).

6. D. Charlesworth, J. H. Willis, Nat Rev Genet. 10, 783–796 (2009).

7. M. Kardos et al., PNAS. 118, e2104642118 (2021).

8. M. Lynch, J. Conery, R. Burger, Am Nat. 146, 489–518 (1995).

9. N. Dussex, H. E. Morales, C. Grossen, L. Dalén, C. van Oosterhout, Trends Ecol Evol. 38, 961–969 (2023).

10. L. F. Keller, D. M. Waller, Trends Ecol Evol. 17, 230–241 (2002).

11. R. Bijlsma, V. Loeschcke, Evol Appl. 5, 117–129 (2012).

12. D. H. Reed, R. Frankham, Conservation Biology. 17, 230–237 (2003).

13. Y. Willi, J. Van Buskirk, A. A. Hoffmann, Annu Rev Ecol Evol Syst. 37, 433–458 (2006).

14. D. A. Bell et al., Trends Ecol Evol, 1–10 (2019).

15. A. R. Whiteley, S. W. Fitzpatrick, W. C. Funk, D. A. Tallmon, Trends Ecol Evol. 30, 42–49 (2015).

16. D. A. Tallmon, G. Luikart, R. S. Waples, Trends Ecol Evol. 19, 489–496 (2004).

17. M. E. Gilpin, M. E. Soulé, in Conservation biology: The science of scarcity and diversity, M. E. Soulé, Ed. (MA: Sinauer Associates, Sunderland, 1986), pp. 19–34.

18. A. R. Weeks et al., Nat Commun. 8, 1071 (2017).

19. C. Vilà et al., Proc R Soc Lond B Biol Sci. 270, 91–97 (2003).

20. W. E. Johnson et al., Science (1979). 329, 1641–1645 (2010).

21. S. W. Fitzpatrick et al., Current Biology, 1–6 (2020).

22. S. M. Miller et al., Conservation Genetics. 21, 41–53 (2020).

23. T. Madsen et al., Current Biology. 30, R1297–R1299 (2020).

24. K. C. Pregler et al., Conserv Lett. 16, e12934 (2023).

25. J. T. Hogg, S. H. Forbes, B. M. Steele, G. Luikart, Proc R Soc Lond B Biol Sci. 273, 1491–1499 (2006).

26. J. G. Clarke, A. C. Smith, C. I. Cullingham, Mol Ecol. 33, e17532 (2024).

27. R. Frankham, Biol Conserv. 195, 33–36 (2016).

28. S. Edmands, Mol Ecol. 16, 463–475 (2007).

29. Z. G. MacDonald et al., Mol Ecol. 34, e17657 (2025).

30. C. C. Kyriazis, R. K. Wayne, K. E. Lohmueller, Evol Lett. 5, 33–47 (2021).

31. S. Nichols et al., Biol Conserv. 290, 110430 (2024).

32. A. Pavlova et al., Evol Appl. 17, 1–20 (2024).

33. P. W. Hedrick, J. A. Robinson, R. O. Peterson, J. A. Vucetich, Anim Conserv. 22, 302–309 (2019).

34. P. W. Hedrick, R. O. Peterson, L. M. Vucetich, J. R. Adams, J. A. Vucetich, Conservation Genetics. 15, 1111–1121 (2014).

35. R. Korona, Evolution (N Y). 53, 1966–1971 (1999).

36. A. S. Kondrashov, D. Houle, Proc R Soc Lond B Biol Sci. 258, 221–227 (1994).

37. K. Ralls et al., Conserv Lett. 11, e12412 (2018).

38. N. Pérez-Pereira, D. Kleinman-Ruiz, A. García-Dorado, H. Quesada, A. Caballero, Mol Ecol, 1–14 (2025).

39. G. West et al., Proc R Soc Lond B Biol Sci. 292, 20242374 (2025).

40. K. Ralls, P. Sunnucks, R. C. Lacy, R. Frankham, Biol Conserv. 251, 108784 (2020).

41. S. Heber, J. V. Briskie, L. A. Apiolaza, PLoS One. 7, e43113 (2012).

42. S. R. Zajitschek, F. Zajitschek, R. C. Brooks, BMC Evol Biol. 9, 1–9 (2009).

43. J. M. Parrett et al., Nat Ecol Evol. 6, 1330–1342 (2022).

44. M. C. Whitlock, A. F. Agrawal, Evolution (N Y). 63, 569–582 (2009).

45. R. J. Dugand, J. L. Tomkins, W. J. Kennington, Nat Commun. 10, 1359 (2019).

46. M. Jarzebowska, J. Radwan, Evolution (N Y). 64, 1283–1289 (2010).

47. J. M. Parrett, R. J. Knell, Proc R Soc Lond B Biol Sci. 285, 20180303 (2018).

48. A. J. Lumley et al., Nature. 522, 470–473 (2015).

49. P. F. Doherty et al., PNAS. 100, 5858–5862 (2003).

50. M. Andersson, *Sexual Selection* (Princeton University Press, New Jersey, 1994).

51. L. Rowe, D. Houle, Proc R Soc Lond B Biol Sci. 263, 1415–1421 (1996).

52. R. Bonduriansky, S. F. Chenoweth, Trends Ecol Evol. 24, 280–288 (2009).

53. A. Plesnar-Bielak, A. M. Skrzynecka, K. Miler, J. Radwan, Evolution (N Y). 68, 2137–2144 (2014).

54. T. Harano, K. Okada, S. Nakayama, T. Miyatake, D. J. Hosken, Current Biology. 20, 2036–2039 (2010).

55. D. Spielman, B. W. Brook, R. Frankham, PNAS. 101, 15261–15264 (2004).

56. M. Lynch et al., Evolution (N Y). 53, 645–663 (1999).

57. J. B. S. Haldane, Am Nat. 71, 337–349 (1937).

58. A. Łukasiewicz, M. Niśkiewicz, J. Radwan, Evolution (N Y). 74, 1851–1855 (2020).

59. R. Joag et al., Genome Biol Evol. 8, 2351–2357 (2016).

60. J. M. Parrett, K. Sobala, S. Chmielewski, K. Przesmycka, J. Radwan, Anim Behav, 123048 (2025).

61. J. Radwan, M. Klimas, Ethol Ecol Evol. 13, 69–79 (2001).

62. A. Plesnar-Bielak, A. M. Skrzynecka, Z. M. Prokop, J. Radwan, Proc R Soc Lond B Biol Sci. 279, 4661–4667 (2012).

63. I. M. Smallegange, Evol Ecol. 25, 857–873 (2011).

64. J. C. Croll, M. Egas, I. M. Smallegange, Journal of Animal Ecology. 88, 11–23 (2019).

65. A. Plesnar-Bielak, A. M. Skwierzyńska, K. Hlebowicz, J. Radwan, BMC Evol Biol. 18, 1–10 (2018).

66. A. Plesnar-Bielak et al., Heredity (Edinb). 133, 45–53 (2024).

67. S. Chmielewski et al., (2024), doi:10.1101/2024.04.15.589577.

68. J. Radwan, M. Czyz, M. Konior, M. Kołodziejczyk, Ethology. 106, 53–62 (2000).

69. A. Plesnar-Bielak, A. Jawor, P. E. Kramarz, Journal of Experimental Biology. 216, 4542–4548 (2013).

70. J. M. Parrett, M. Kulczak, N. Szudarek-Trepto, Oikos, 1–10 (2024).

71. J. Radwan, Heredity (Edinb). 74, 669–673 (1995).

72. J. M. Parrett et al., Evolution (N Y). 77, 1289–1302 (2023).

73. F. T. Rhebergen, K. A. Stewart, I. M. Smallegange, Ecol Evol. 12, 1–9 (2022).

74. A. Łukasiewicz, N. Porwal, M. Niśkiewicz, J. M. Parrett, J. Radwan, Evolution (N Y). 77, 2291–2300 (2023).

75. J. Radwan, Heredity (Edinb). 90, 371–376 (2003).

76. A. M. Bolger, M. Lohse, B. Usadel, Bioinformatics. 30, 2114–2120 (2014).

77. H. Li, (2013) (available at http://arxiv.org/abs/1303.3997).

78. H. Li et al., Bioinformatics. 25, 2078–2079 (2009).

79. R. Kofler, R. V. Pandey, C. Schlötterer, Bioinformatics. 27, 3435–3436 (2011).

80. R. Kofler et al., PLoS One. 6, e15925 (2011).

81. H. Li, Bioinformatics. 27, 2987–2993 (2011).

82. A. R. Quinlan, I. M. Hall, Bioinformatics. 26, 841–842 (2010).

83. P. Cingolani et al., Fly (Austin). 6, 80–92 (2012).

84. R Development Core Team, R: A language and environment for statistical computing. (2024), (available at https://www.r-project.org/).

85. H. Wickham, ggplot2: Elegant Graphics for Data Analysis (2016).

86. M. E. Brooks et al., R Journal. 9, 378–400 (2017).

87. T. M. Therneau, coxme: Mixed Effects Cox Models (2020), (available at https://cran.r-project.org/web/packages/coxme/).

